# Chromosome-scale genome of the woody oilseed crop sacha inchi elucidates the molecular basis of alpha-linolenic acid biosynthesis and triacylglycerol accumulation in seeds

**DOI:** 10.64898/2026.03.18.712556

**Authors:** Bang-Zhen Pan, Xuan Zhang, Xiao-Di Hu, Qiantang Fu, Mao-Sheng Chen, Yan-Bin Tao, Long-Jian Niu, Huiying He, Yi Shen, Zhukuan Cheng, Tiange Lang, Changning Liu, Zeng-Fu Xu

## Abstract

Sacha inchi (Plukenetia volubilis L.) is an emerging woody oilseed crop prized for its high alpha-linolenic acid (ALA) content. Despite its nutritional and economic value, the lack of high-quality genomic resources has hindered genetic improvement and the elucidation of its unique polyunsaturated fatty acid and lipid biosynthetic pathways. In this study, we report a high-quality, chromosome-scale genome assembly of sacha inchi with a total length of 710.62 Mb, integrated from Illumina, PacBio, and chromosome conformation capture (Hi-C) technology. The genome harbors 37,570 protein-coding genes, and 379.86 Mb (53.45%) of repetitive sequences. Phylogenomic analysis reveals that sacha inchi diverged from its closest relative Ricinus communis, ∼ approximately 36.2 million years ago. Comparative genomics indicates that sacha inchi experienced only ancient whole genome duplication events. To elucidate the mechanisms governing ALA biosynthesis and triacylglycerol (TAG) accumulation in sacha inchi seeds, we performed temporal transcriptome profiling across six seed development stages. Our findings demonstrate that high TAG content is primarily driven by the sustained expression of biosynthetic genes and low activity of degradation genes during mid-to-late seed development. Notably, while genes encoding stearoyl-ACP desaturases (SADs) maintain the precursor pool, the expression of genes encoding fatty-acid desaturase 2 (FAD2) and fatty-acid desaturase 3 (FAD3) is positively correlated with the final accumulation of C18:2 and C18:3 fatty acids. We also identified lncRNAs as potential epigenetic regulators of these key pathways. This high-quality genome provides a critical foundation for elucidating the molecular mechanisms of seed growth and development in sacha inchi.

## INTRODUCTION

*Plukenetia volubilis* L., also known as sacha inchi and Inca peanut, is native to the high-altitude rain forests of the Andean region of South America, and is a member of the Euphorbiaceae family (Hamaker et al., 1992). Sacha inchi, as shown in Fig. S1, grows as a woody monoecious vine bearing slender racemose inflorescences, with one or two pistillate flowers single at the basal node and numerous staminate flowers in condensed cymes at the distal nodes, yielding a star-shaped fruit capsule with 4-7 lobes, each containing one oleaginous seed (Gillespie, 1993, Gillespie, 2007, Fu et al., 2014). The seeds have been a daily food component of the indigenous people in South America (Hamaker et al., 1992). This plant has been extensively investigated for its potential to yield nutritional, cosmetic and pharmaceutical products (Goyal et al., 2022, Abd Rahman et al., 2023). Sacha inchi seeds contain 35–60% lipids, 25–30% proteins, tocopherols, polyphenols, and sterols (Niu et al., 2014, Hamaker et al., 1992, Bondioli et al., 2006, Chirinos et al., 2013, Valencia et al., 2024). The important roles of C18:3 fatty acids (C18:3 FAs) (Simopoulos, 1999) and the high ratio of C18:3 FAs to C18:2 FAs in human health have been previously recognized (Simopoulos, 1991). The seed oil of sacha inchi contains a high level (40∼50%) of alpha-linolenic acid (ALA), a type of C18:3 FA, and a relatively low proportion (∼30%) of linoleic acid (LA), a type of C18:2 FA (Bondioli et al., 2006, Wang and Liu, 2014, Fu et al., 2024a); thus, this seed oil could help people meet the requirements for increasing the ratio of C18:3 FAs to C18:2 FAs in their daily diets. The expanding cultivation of sacha inchi beyond its native range for commercial purposes has intensified breeding efforts aimed at improving agronomic traits, such as seed yield while preserving its prized oil profile (Supriyanto et al., 2022, Istiandari and Faizal, 2025). Molecular breeding strategies for such optimization, however, are currently hampered by the lack of comprehensive genomic resources. A foundational step toward this goal is to elucidate the genetic determinants of its hallmark trait—the high seed oil enriched in ALA.

Although most pathways of FA and triacylglycerol (TAG) biosynthesis have been identified in *Arabidopsis thaliana* (Li-Beisson et al., 2013) and some oil crops (Lin et al., 2022, Chen et al., 2019, Sreedhar et al., 2015, Unver et al., 2017, Zuo et al., 2024), only a limited number of transcriptomic studies in sacha inchi have been reported (Wang et al., 2012, Wang and Liu, 2014, Hu et al., 2018, Fu et al., 2022, Liu et al., 2020). These studies provide preliminary genetic insights into oil production. Furthermore, recent methodological advances, such as the development of *Agrobacterium*-mediated genetic transformation systems in sacha inchi (Lin et al., 2025, Yu et al., 2025), have provided valuable tools for in planta gene functional validation. However, the successful application of these tools for elucidating complex traits, such as oil biosynthesis, relies on the prior identification of candidate genes, a process that is severely hindered by the absence of a high-quality, chromosome-scale reference genome of sacha inchi.

In this study, to unveil the genetic basis of the unique composition of seed oil in sacha inchi, particularly the accumulation of high levels of ALA, a combination of second- and third-generation sequencing technologies (Illumina sequencing and Pacific Biosciences sequencing, respectively), and chromosome conformation capture (Hi-C) technology were employed to sequence and obtain a chromosome-scale assembly of sacha inchi genome. Through detailed genome annotation and transcriptome analysis of seeds at six different developmental stages, we identified key genes and gene families involved in the FA and TAG biosynthesis pathways. This genome sequence not only provides crucial insights into the molecular mechanisms underlying high ALA production in sacha inchi, but also offers a valuable genomic resource for functional genomics and molecular breeding research on this important woody oilseed crop.

## RESULTS

### Genome sequencing and assembly

A one-year-old sacha inchi plant was selected for whole genome sequencing. To facilitate the assembly of the genome, the chromosome number of sacha inchi was first analyzed. The results showed that the chromosome number of sacha inchi is 2x = 58 (Fig. 1A), which is in line with a previous finding that, among 72 cells observed in sacha inchi, the most common chromosome number is 2x = 58 (Cai et al., 2013).

**Fig. 1.**
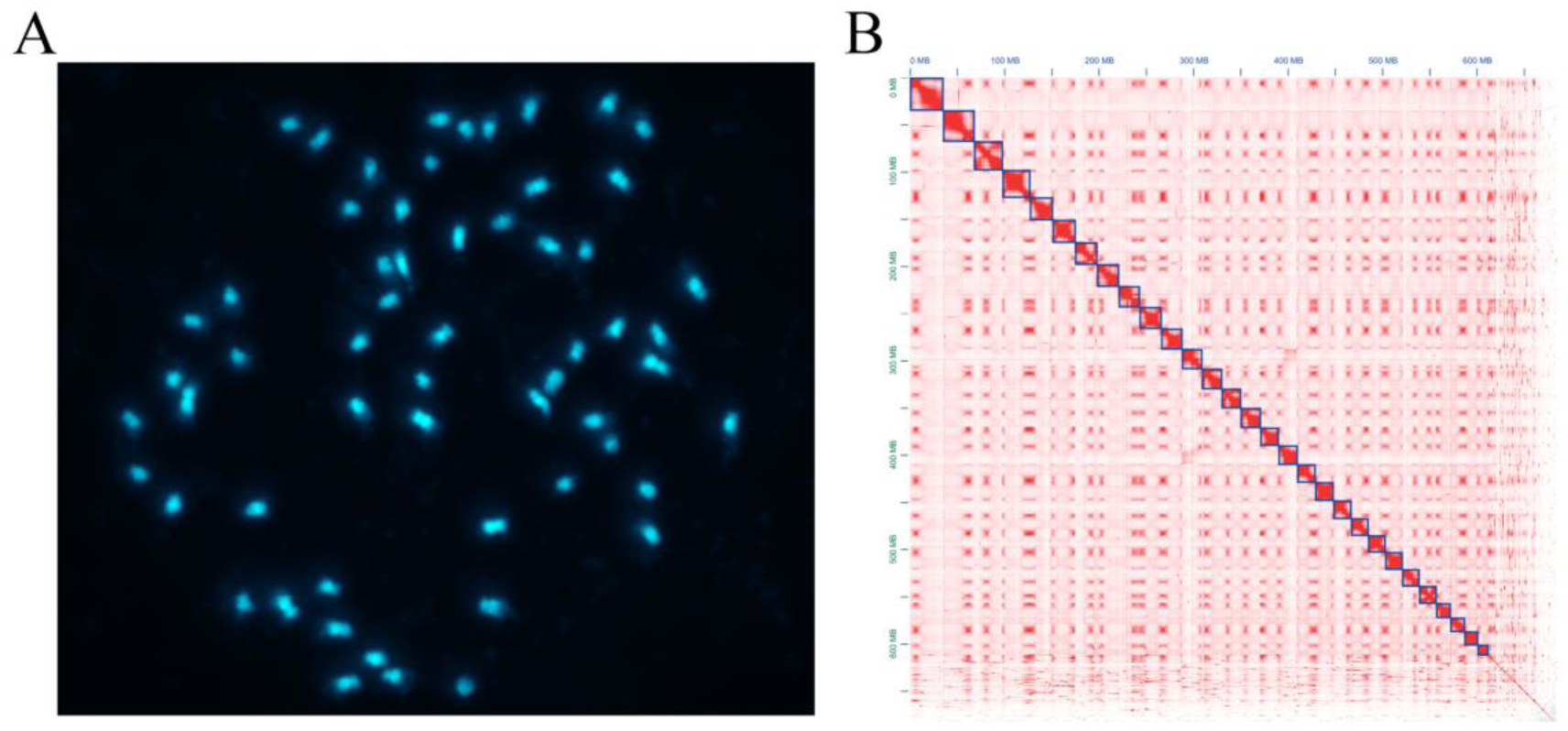
Chromosome number and chromosome-scale assembly of the sacha inchi genome (A) Chromosomes stained with DAPI. Bar = 5 μm. The chromosome number of sacha inchi is 2x = 58. (B) Genome wide Hi-C heat map.

Using the Illumina HiSeq 2500 platform, a total of 275.34 Gb of clean paired-end reads were generated, corresponding to more than 397 X coverage of the sacha inchi genome, with an estimated size of 689.37 Mb based on K-mer analysis (Supplemental Table S1 and Supplemental Fig. S2). *De novo* assembly of these sequences with Allpaths-LG software (Gnerre et al., 2011) yielded a 707.39 Mb assembly with a contig N50 of 82.55 Kb and a scaffold N50 of 1.32 Mb (Table 1). For further gap closing, long read sequencing was conducted via PacBio sequencing (Supplemental Tables S2 and S3), which yielded a 716.52 Mb assembly with a contig N50 of 194.13 Kb and a scaffold N50 of 1.34 Mb (Table 1). Hi-C technology was then used to anchor the assembled sequences into chromosomes, which anchored 86.32% of the assembled sequences into 29 chromosomes, with maximum and minimum lengths of 35.98 and 11.04 Mb, respectively (Fig. 1B, Supplemental Table 4). The final high-quality assembly of 710.62 Mb comprised 29 chromosomes and 14504 scaffolds with an N50 of 20.37 Mb (Table 1), which is close to the genome size estimated from K-mer analysis and flow cytometry (Supplemental Figs. S2 and S4). The GC content of the sacha inchi genome was 29.97% (Table 2, Supplemental Fig. S5), which is lower than that of most plant species whose genomes have been sequenced (Singh et al., 2013, Singh et al., 2016). Compared with the published genomes of other spurge species, the GC content in the intergenic regions of sacha inchi genome is remarkably low, whereas the GC contents in genes and exons are similar (Cui et al., 2018, Chan et al., 2010, Tang et al., 2016, Wu et al., 2015).

**Table 1.** Statistic summary of sacha inchi genome assembly.

| Category | Number | Size<br>(Mb) | N50<br>(Kb) | Max<br>Length<br>(Mb) | Gap<br>(%) |
| --- | --- | --- | --- | --- | --- |
| Estimate of genome size (flow cytometry) | - | 691.93 | - | - | - |
| Estimate of genome size (K-mer) | - | 689.37 | - | - | - |
| Contig (HiSeq) | 29,791 | 670.08 | 82.55 | 1.02 | - |
| Contig (HiSeq+PacBio) | 23,565 | 689.93 | 194.13 | 1.91 | - |
| Contig (HiSeq+PacBio+HiC) | 26,276 | 689.93 | 153.14 | 1.38 | - |
| Scaffold (HiSeq) | 13,818 | 707.39 | 1323.50 | 8.79 | 5.27 |
| Scaffold (HiSeq+PacBio) | 13,816 | 716.52 | 1339.21 | 8.84 | 3.71 |
| Scaffold (HiSeq+PacBio+HiC) | 14,533 | 710.62 | 20370.02 | 35.98 | 3.83 |

**Table 2.**
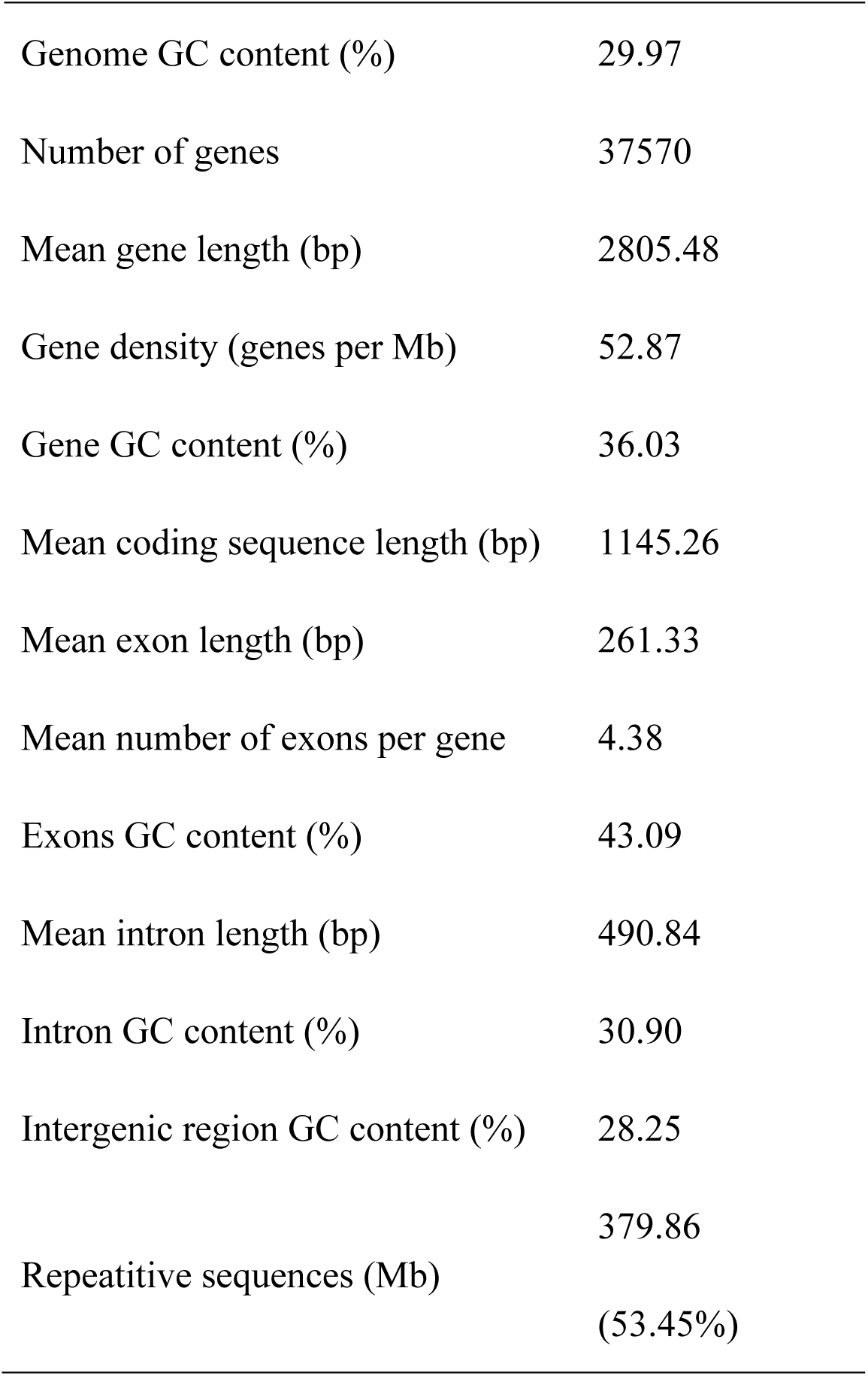
Statistics of sacha inchi genome annotation.

To evaluate the assembly quality, Illumina reads (27.24 Gb of data, ∼35.26 X) from the 220 bp library were mapped to the assembled genome with BWA (Li and Durbin, 2010). The read mapping rate and coverage of the genome were 95.24% and 89.50%, respectively, indicating a low frequency of misassembly in the genome (Supplemental Table S5). The completeness of the genome was checked by mapping 1440 Benchmarking Universal Single-Copy Orthologs (BUSCOs) to the assembled genome using BUSCO v3.0.2 (Simao et al., 2015); this verification revealed that 94.3% of the BUSCOs were covered by the assembled genome, of which 74.2% were single, 20.1% were duplicated, and only 2.2% had fragmented matches (Supplemental Table S6). Moreover, weighted score of highly conserved core gene families (coreGFs) (Van Bel et al., 2012) for sacha inchi was 98.6% (Supplemental Table S6). The results of these evaluations demonstrated that the assembly of sacha inchi genome was of high quality.

### Genome annotation

The sacha inchi genome contains a total of 379.86 Mb (53.45%) of repetitive sequences (Table 2, Supplemental Table S7), which is similar to that of *R. communis* (50%) (Chan et al., 2010) and *Mercurialis annua* (51.5%) (Veltsos et al., 2019). The most abundant transposable element (TE) is the long terminal repeat (LTR) family, accounting for ∼46% of the genome assembly (Supplemental Table S8). We investigated the retrotransposon activities in sacha inchi by dating the insertion time of all complete LTR structures based on divergence via distances of 5’ and 3’ solo-LTRs, and found that the insertion time of LTRs peaked approximately 1.6 million years ago (Mya) (Supplemental Fig. S6).

Protein-coding genes were predicted based on a combination of *ab initio*, conserved protein homologs and assembled transcripts that were recruited from the transcriptomes of six developmental stage seeds from 1 week to 17 weeks (mature) after pollination (WAP), which we generated (Supplemental Fig. S7), and other published transcripts of different tissues (Hu et al., 2018, Fu et al., 2018, Wang et al., 2012). We predicted genes from repeat-masked genome sequences. All the results from the three types of predictions were integrated by GLEAN (Elsik et al., 2007). As a result, we predicted a total of 37,570 protein-coding genes with average lengths of 1145.3 bp, 261.3 bp and 490.8 bp for coding sequences, exons and introns, respectively (Table 2, Supplemental Table S9). Genome comparisons between sacha inchi and seven other plant species showed differences in gene numbers, average lengths of mRNAs/coding-sequences/introns/exons, average numbers of exons per gene, and proportions of genomic GC content (Supplemental Fig. S8 and Supplemental Table S10). The BUSCO evaluation demonstrated that 93% and 4% of the 1440 BUSCOs were complete and fragmented, respectively (Supplemental Table S11).

Annotation of the predicted genes was performed by aligning their amino acid sequences against a number of protein databases, including InterPro (Burge et al., 2012), KEGG (Kanehisa and Goto, 2000), Gene Ontology (Ashburner et al., 2000), Swiss-Prot (Bairoch and Apweiler, 2000), NR and TrEMBL (Boeckmann et al., 2003). Overall, a total of 33,959 (90.39%) of the protein-coding genes could be annotated by at least one of the databases above (Supplemental Table S12). In addition, non-coding RNAs were identified by searching against various RNA libraries. We ultimately identified 129 miRNAs, 158 rRNAs, 543 tRNAs,

### Comparative phylogenomic and whole-genome duplication analysis of sacha inchi and related plant

To clarify the evolutionary position and genomic evolutionary characteristics of sacha inchi, we performed gene family clustering, phylogenetic analysis, and whole-genome duplication (WGD) analysis using 13 representative plant species, including six spurge plant species (sacha inchi, *Ricinus communis*, *Jatropha curcas*, *Manihot esculenta*, *Hevea brasiliensis* and *Vernicia fordii*), *Actinidia chinensis*, *Arabidopsis thaliana*, *Elaeis guineensis*, *Linum usitatissimum*, *Populus trichocarpa*, *Vitis vinifera* and *Oryza sativa*. Clustering homologs of all 13 species, revealed a total of 28,430 gene families, among which 6,469 gene families were common to all 13 species, whereas 556 were confined to eudicots (all clustered species except *O. sativa*). In the sacha inchi genome, 31,483 genes were assigned to 15,108 families, including 362 unique families (Supplemental Fig. S9 and Supplemental Table S14). Comparative analysis with other five spurge species (*R. communis*, *J. curcas*, *M. esculenta*, *H. brasiliensis* and *V. fordii*), revealed that 11,418 gene families were conserved across all six species, while 628 families were unique to sacha inchi (Fig. 2A).

**Fig. 2.**
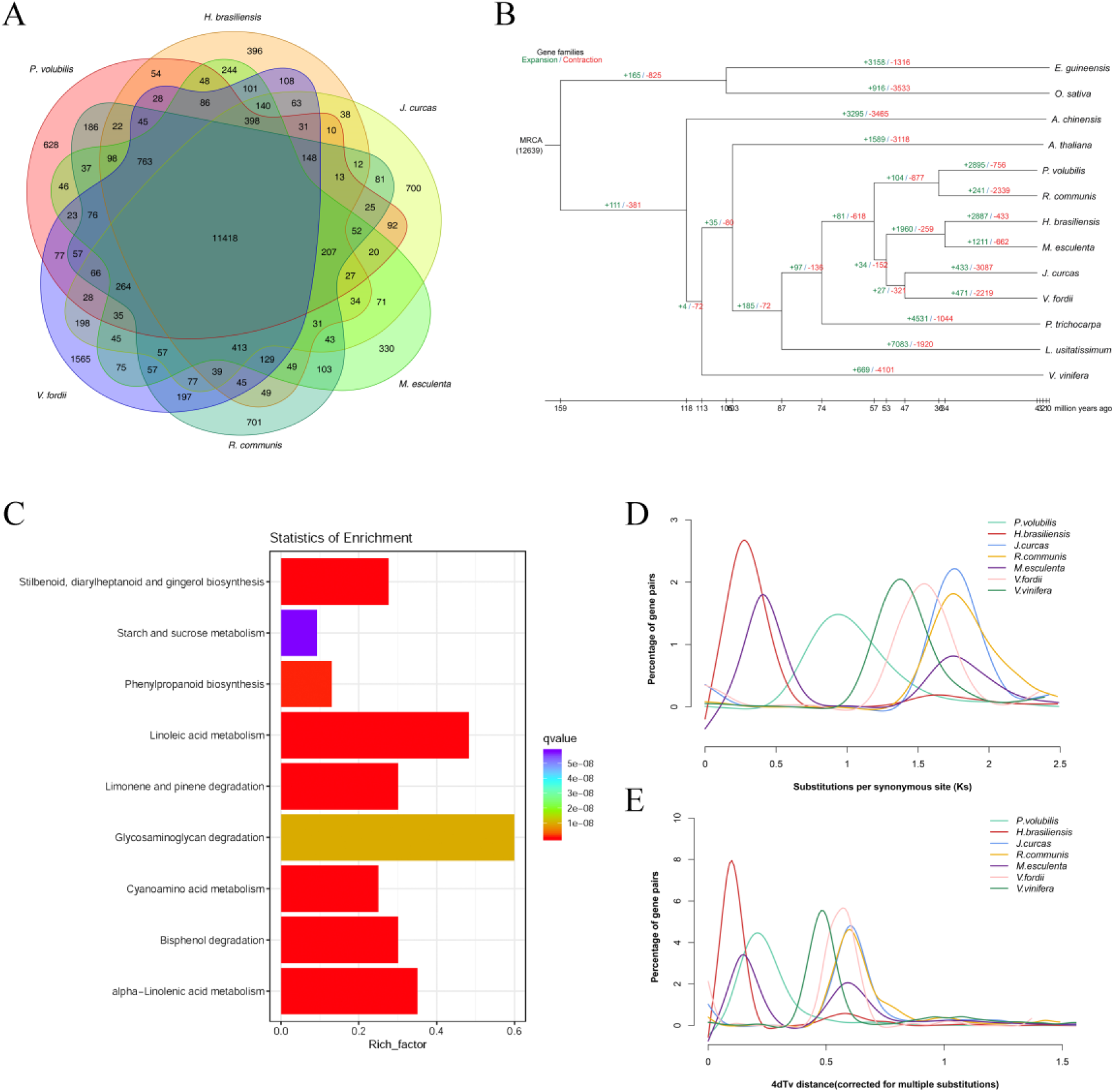
Genome evolution analysis of sacha inchi. (A) Venn diagram of orthologous gene families of sacha inchi and other Euphorbiaceae plants. (B) Phylogeny and divergence time estimation by molecular clock analysis. The number at the root represents the divergence time. The values above each branch denote the gene family expansion/contraction number at each round of genome duplication after diversifying from the common ancestor. (C) KEGG enrichment of the expanded gene families (*p*<0.05). (D) and (E) Density distributions of Ks and 4DTv, respectively, for paralogous genes of the sacha inchi genome and other selected species.

A phylogenetic tree of the 13 species was constructed using 1:1 single-copy gene families. Proteins and CDSs from single-copy families were aligned with MUSCLE (3.8.31) (Edgar, 2004a), and four-fold degenerate sites were extracted and merged into a supergene as an input of MrBayes (Huelsenbeck and Ronquist, 2001) with *O. sativa* as the outgroup. The results estimated the divergence time of the Euphorbiaceae family at ∼54.4 Mya (Fig. 2B), consistent with a previously reported date of ∼58 Mya (Tang et al., 2016). Sacha inchi was found to be most closely related to *R. communis*, with a common ancestor dating back *∼*32.6 Mya (Fig. 2B).

Gene family expansion and contraction were analyzed using the CAFÉ program (De Bie et al., 2006) by comparing family size differences between the most recent common ancestor and each of the 13 plant genomes. Since splitting from its most recent common ancestor, sacha inchi has experienced 2895 gene family expansion events and 756 contraction events (Fig. 2B). Using conditional likelihoods as the test statistics with a significance threshold of *P* < 0.05, we identified significantly expanded gene families in sacha inchi. KEGG enrichment analysis of these families revealed significant associations with starch and sucrose metabolism, linoleic acid metabolism, and alpha-linolenic acid (ALA) metabolism (Fig. 2C). Notably, genes annotated to the latter two lipid metabolism pathways were predominantly lipoxygenases (LOXs) and 12-oxophytodienoate reductases (OPRs), which are involved in the oxidative branch leading to jasmonic acid (JA) biosynthesis (Viswanath et al., 2020, Yi et al., 2024). As the expression of these JA biosynthetic genes generally decreased during the middle and late stages of seed development (Supplemental Table S15; Supplemental Fig. S10), their family expansion likely reflects an evolutionary adaptation to other selective pressures, such as enhanced defense against biotic stresses (e.g., pests and pathogens), rather than a direct role in driving high ALA accumulation in sacha inchi seeds.

To further explore the evolutionary mechanisms underlying these genomic features, we conducted WGD analysis by comparing synonymous substitution rates (Ks) and fourfold synonymous third-codon transversion (4DTv) distributions among paralogues and orthologues of six Euphorbiaceae species and *V. vinifera*. The Ks and 4DTv distributions of sacha inchi are similar to those of *J. curcas*, *R. communis*, and *V. fordii*, which display only one obvious peak (Fig. 2D, E), indicating a single WGD event during the genome evolution of these four species (Chan et al., 2010, Tang et al., 2016, Cui et al., 2018, Ha et al., 2019). This result was supported by syntenic analysis between sacha inchi and *J. curcas*, and between sacha inchi and *M. esculenta* (Supplemental Figs. S11 and S12). Combined with previous reports, our analysis revealed that all six Euphorbiaceae species shared a common ancient WGD event, whereas only *M. esculenta* and *H. brasiliensis* experienced an additional recent WGD event (Fig. 2D, E) (Chan et al., 2010, Tang et al., 2016).

### Differential expression of oil-related genes during seed development revealed strategies for oil accumulation in sacha inchi

To understand the molecular mechanisms underlying these characteristics of sacha inchi seed oil, we analyzed key genes in the FA and TAG biosynthesis pathways across different stages of seed development, ranging from 1 to 17 WAP (Supplemental Fig. S7). According to Bates et al.(2013) and Vanhercke et al.(2019), lipid biosynthesis was categorized into four metabolic modules: *Push*, *Pull*, *Package* and *Protect*, whereas *Push* refers to *de novo* FA biosynthesis in plastids, *Pull* refers to neutral TAG assembly through the Kennedy pathway in the endoplasmic reticulum, *Package* refers to lipid droplet (LD) biosynthesis in the cytosol, and *Protect* refers to the blockage of TAG turnover through the downregulation of lipases and/or key genes in the β-oxidation pathway in the peroxisome (Fig. 3).

**Fig. 3.**
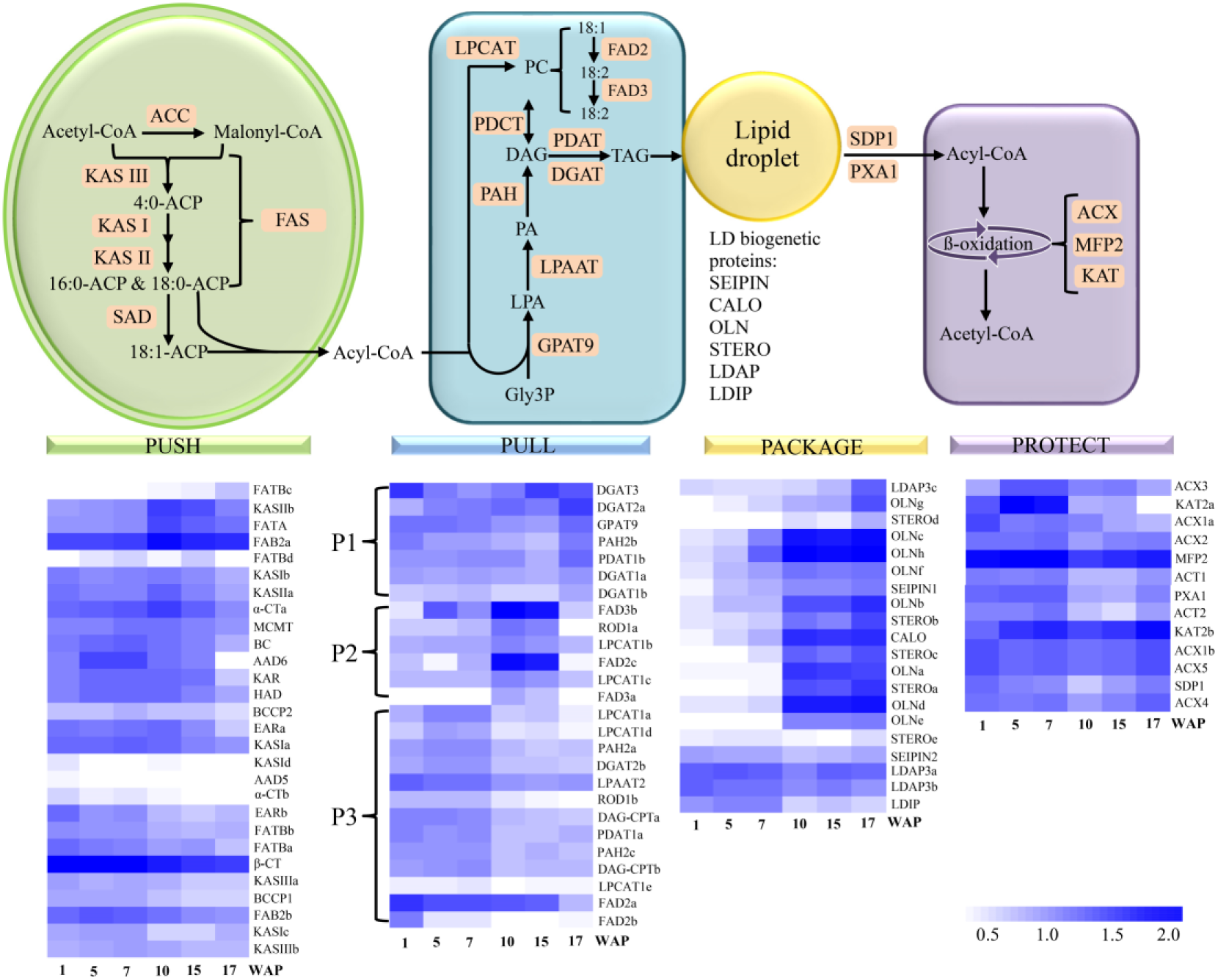
Expression profiles of oil biosynthesis-related genes in sacha inchi seeds. The color scale at the bottom shows log_10_ (RPKM+1) values. In the *Push* module, *BCCP1, Biotin Carboxyl Carrier Protein of Heteromeric ACCase 1; BCCP2, Biotin Carboxyl Carrier Protein of Heteromeric ACCase 2; BC, Biotin Carboxylase of Heteromeric ACCase; α-CT, Carboxyltransferase alpha Subunit of Heteromeric ACCase; β-CT, Carboxyltransferase beta Subunit of Heteromeric ACCase; MCMT, Malonyl-CoA: ACP Malonyltransferase; KAR, Ketoacyl-ACP Reductase; EARa/b, Enoyl-ACP Reductase a/b; HAD, Hydroxyacyl-ACP Dehydrase; KASI, Ketoacyl-ACP Synthase I; KASII, Ketoacyl-ACP Synthase II; KASIII, Ketoacyl-ACP Synthase III; FATB, Acyl-ACP Thioesterase B; FATA, Acyl-ACP Thioesterase A; FAB2, Fatty Acid Biosynthesis 2; DES5, Stearoyl-acyl Carrier Protein-desaturase 5; DES6, Stearoyl-acyl Carrier Protein-desaturase 6*. In the *Pull* module, P1, expression pattern 1; P2, expression pattern 2; P3, expression pattern 3. *GPAT9, Glycerol-3-Phosphate Acyltransferase; LPAAT2, Lysophosphatidic Acid Acyltransferase 2; PAH2, Phosphatidate Phosphatase 2; ROD1, reduced oleate desaturation 1; DAG-CPT, Diacylglycerol Cholinephosphotransferase; LPCAT1, 1-Acylglycerol-3-Phosphocholine Acyltransferase Lysophospholipid acyltransferase; DGAT1, Acyl-CoA: Diacylglycerol Acyltransferase 1; DGAT2, Acyl-CoA: Diacylglycerol Acyltransferase 2; DGAT3, Acyl-CoA: Diacylglycerol Acyltransferase 3; PDAT1, Phospholipid: Diacylglycerol Acyltransferase 1; FAD2, Fatty acid desaturases 2; FAD3, Fatty acid desaturases 3.* In the *Package* module, *CALO, Caleosin; OLN, Oil-Body Oleosin; STERO, Steroleosin; LDAP3, LD-associated protein 3; LDIP, LDAP-interacting protein; SEIPIN1; SEIPIN2.* In the *Protect* module, *SDP1, Sugar dependant 1; PXA1, Peroxisomal ABC Transporter; ACT1, acyl-CoA thioesterase 1; ACT2, acyl-CoA thioesterase 2; ACX1, acyl-CoA oxidase 1; ACX2, acyl-CoA oxidase 2; ACX3, acyl-CoA oxidase 3; ACX4, acyl-CoA oxidase 4; ACX5, acyl-CoA oxidase 5; MFP2, multifunctional protein 2; KAT2, 3-ketothiolase 2*.

The results revealed that most of the genes in the *Push* module were expressed consistently during seed development, with a certain degree of reduced expression in the seeds at 17 WAP (Fig. 3, Supplemental Table S16), indicating that FA biosynthesis began in the very early stage and occurred throughout seed development, most of which exist in the form of free FAs and phospholipids in seeds at early stages (Supplemental Fig. S13) (Wang and Liu, 2014). *Fatty acid biosynthesis 2a* and *2b* (*FAB2a* and *FAB2b*), and *stearoyl-acyl carrier protein-desaturase* (*AAD6*), which encode stearoyl-ACP desaturases (SADs) and catalyze the production of C18:1 fatty acids (Kazaz et al., 2020), were predominantly expressed during seed development. In the *Pull* module, there are three main gene expression patterns. The first pattern includes genes encoding glycerol-3-phosphate acyltransferase 9 (*GPAT9*), phosphatidate phosphohydrolases (*PAH2b*), diacylglycerol acyltransferase (*DGAT1a, DGAT1b DGTA2a* and *DGTA3*), and phospholipid:diacylglycerol acyltransferase (*PDAT1b*), most of which are responsible for the first and last steps of the Kennedy pathway. These genes were expressed throughout the six development stages, echoing the continuous biosynthesis of FAs in the *Push* module. The second pattern includes genes encoding lysophosphatidylcholine acyltransferase (*LPCAT1b* and *LPCAT1c*), fatty-acid desaturase (*FAD2c*, *FAD3a,* and *FAD3b*), and phosphatidylcholine: diacylglycerol cholinephosphotransferase (*reduced oleate desaturation 1a*, *ROD1a*), which are the critical steps involved in TAG biosynthesis via phosphatidylcholine (PC) editing. These genes were expressed predominantly in the seeds from 5 to 15 WAP and peaked at 10 and 15 WAP (Fig. 3, Supplemental Table S16), indicating that the unsaturation degree of the final TAG was determined mainly in these two stages, which was supported by the increased contents of C18:2 and C18:3 FAs at the corresponding stages (Supplemental Fig. S13) (Wang and Liu, 2014). For the third pattern, genes including *PAH2a, PAH2c, ROD1b, LPCAT1a, LPCAT1d, LPCAT1e*, *FAD2a, FAD2b, DGTA2b, PDAT1a* and the genes encoding lysophosphatidic acid acyltransferase 2 (*LPAAT2*) and CDP-choline:diacylglycerol cholinephosphotransferase (*DAG-CPTa, DAG-CPTb*), were more abundant in the seeds at 1, 5 and 7 WAP, suggesting that part of the TAG was assembled during the early development of the seeds.

A previous report revealed that early generated TAG consists mainly of saturated fatty acids (Wang and Liu, 2014). Most of the genes encoding LD biogenetic proteins in the *Package* module display an obvious low-high expression pattern. They had lower abundance in the seeds at 1, 5 and 7 WAP, compared with those at 10, 15 and 17 WAP, indicating that the seeds produced LD primarily in the late development stages. This result was also confirmed by the high level of TAG (neutral lipids) at the middle and late development stages (Supplemental Fig. S13) (Wang and Liu, 2014). Most plants have three *SEIPIN* genes, which control the size of the LD, with SEIPIN1 producing a large LD, SEIPIN2 producing a normal size of the LD, and SEIPIN3 producing a small LD (Cai et al., 2015). We identified one gene encoding SEIPIN1 and three genes encoding SEIPIN2 in the sacha inchi genome. Like most other genes that encode LD biogenetic proteins, the gene encoding SEIPIN1 was expressed predominantly in seeds at 10, 15 and 17 WAP. Only one of three genes encoding SEIPIN2 was expressed consistently during seed development. These results indicate that the production of normally sized LDs occurred throughout seed development, whereas the production of large LDs happened in the middle and late stages.

The *Protect* module basically comprises TAG lipases, peroxisomal ABC transporter 1 (PXA1), and elements involved in the β-oxidation pathway, most of which showed a high-low expression pattern. They had higher abundance in the seeds at 1, 5 and 7 WAP, compared with those at 10, 15 and 17 WAP, indicating that at middle and late development stages, the metabolism rate of TAG was lower than that at early stages, and thus resulting in rapid accumulation of TAG. In addition, the expression profiles of other TAG lipases and genes involved in the ALA metabolism pathway identified in the sacha inchi genome also support this result (Supplemental Figs. S14 and S15). Taken together, our results show that the ultimate TAG content was governed by the consistent expression of genes involved in FA biosynthesis and TAG synthesis during seed development, and the relatively low expression of genes involved in TAG degradation in the middle and late stages.

### The expression profiles of *FAD2* and *FAD3* reflect the contents of C18:2 and C18:3 FAs

Given that polyunsaturated FAs have been proven to benefit human health, particularly because they have positive effects on the control of cardiovascular disease (Simopoulos, 1991), oil composition determines the value and utility of seed oil. Therefore, what affects the degree of unsaturation of TAG? Here, we analyzed the relationships between the mRNA abundance of genes encoding SAD, FAD2 and FAD3, and the content of C18:2 and C18:3 FAs in the seed oils of sacha inchi, *P. frutescens*, and *L. usitatissimum* (C18:3 FA content > 45%), *A. thaliana* (C18:3 FA content ∼20%), and *A. hypogaea*, *G. hirsutum* and *J. curcas* (C18:3 FA content < 1%), which were categorized into high, middle and low C18:3 FA content groups, respectively. First, we found that the expression levels of genes encoding SAD, FAD2 and FAD3 during seed development displayed similar patterns among middle and high C18:3 FA-containing oilseed crops, peaking at the middle-late development stages, whereas the lowest values occurred at the early and late (and/or mature) stages (Fig. 4A-D). Second, in low C18:3 FA-containing oilseed crops (*A. hypogaea*, *G. hirsutum* and *J. curcas*), genes encoding SADs maintain a high level of expression throughout seed development, whereas *FAD3* is slightly expressed in the early stage and shut down in later stages, which may imply little accumulation of C18:3 FA (Fig. 4E-G).

**Fig. 4.**
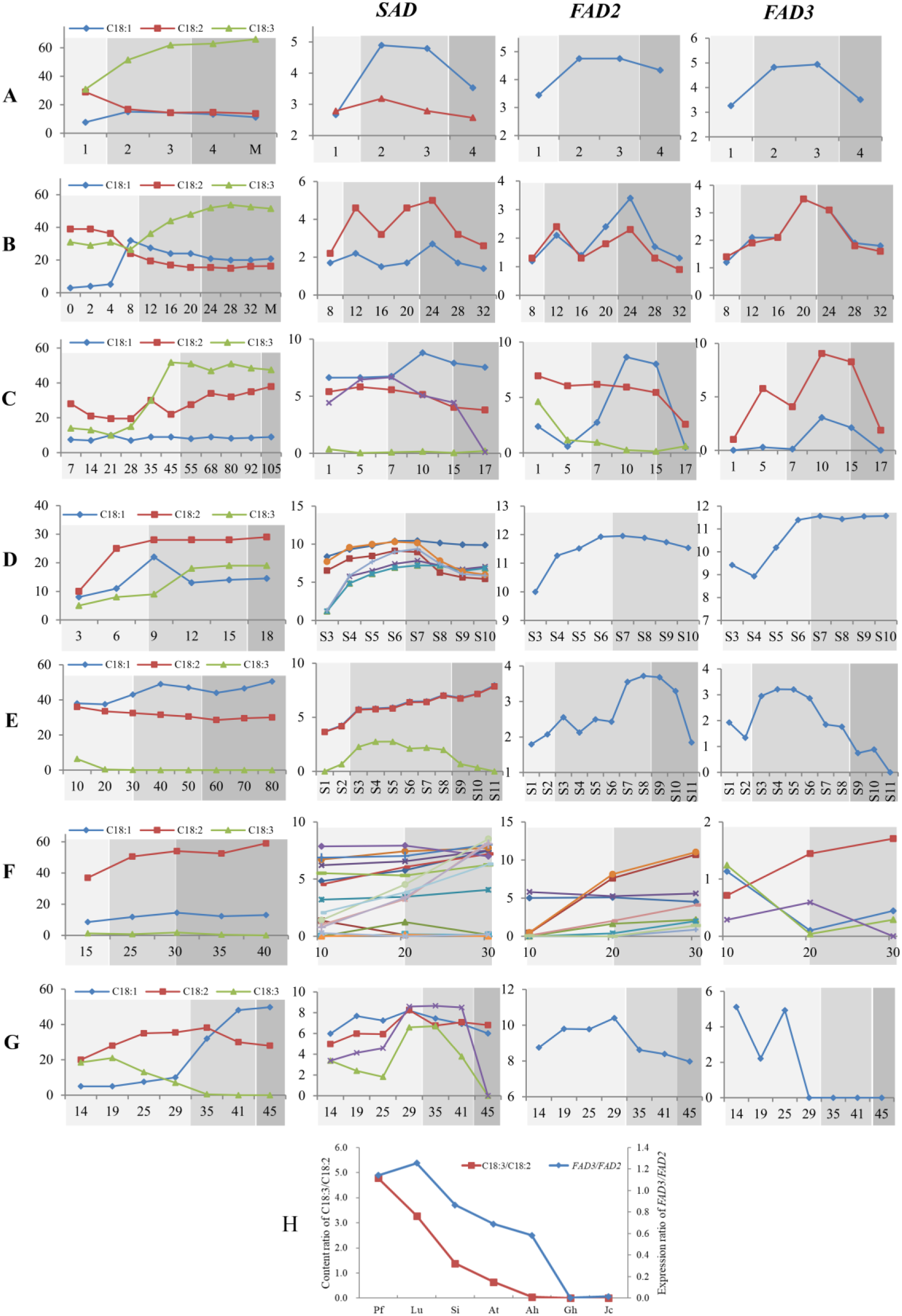
Relationships between the contents of C18:1, C18:2 and C18:3 FAs and the expression levels of genes encoding SAD, FAD2 and FAD3. (**A)** *P. frutescens*. (**B)** *L. usitatissimum*. **(C)** Ssacha inchi. (**D)** *A. thaliana.* **(E)** *A. hypogaea*. **(F)** *G. hirsutum* (**G)** *J. curcas*. (**H)** Positive correlation between the *FAD3/FAD2* expression ratios and the C18:3 and C18:2 content ratios in oilseed crops. Pf, *P. frutescens*. Lu, *L. usitatissimum*. Si, Ssacha inchi. At, *A. thaliana.* Ah, *A. hypogaea*. Gh, *G. hirsutum.* Jc, *J. curcas*. The details of the data in this figure are listed in Supplemental Table S17. Lines with different colors in the figures with different expression levels represent different copies of genes. Varying shading intensities (from light to dark) represent seeds at three developmental stages: the early stage (prior to rapid lipid accumulation), the middle stage (rapid lipid accumulation), and the late stage (plateau phase).

Regarding *FAD2*, its expression was also positively correlated with the content of C18:2 FA. For *A. hypogaea* and *J. curcas* (containing ∼20% C18:2 FA), *FAD2* was moderately expressed (lower than *SAD*) in both the early and late stages. In *G. hirsutum*, in addition to more genes encoding FAD2, these genes were alternatively expressed at high levels across the three selected developmental stages (early and middle stages), which is consistent with the high content of C18:2 FA (∼59%). Moreover, the content ratios of C18:2 and C18:3 FAs of these species match well with the expression ratios of *FAD2* and *FAD3* (Fig. 4H). Consequently, the data indicated that the expression levels of *FAD2* and *FAD3* are positively correlated with the contents of C18:2 and C18:3 FAs, respectively. The *FAD2*/*FAD3* expression ratio may be an important factor that regulates the relative contents of C18:2 and C18:3 FAs.

### Identification of candidate ncRNAs associated with high contents of total oil and unsaturated FAs

In addition to protein-coding RNAs, gene regulation by non-coding RNAs (ncRNAs) is also essential for various aspects of plant development. Here, the transcriptomes of ncRNAs, including microRNAs (miRNAs), circular RNAs (circRNAs), and long non-coding RNAs (lncRNAs) were analyzed to characterize the functions of candidates in regulating oil-related genes.

MiRNAs act as post-transcriptional regulators of gene expression by targeting mRNAs for cleavage or translational repression (Voinnet, 2009). Our results revealed that 158 miRNAs were expressed at least at one stage during seed development, with more read counts distributed in the seeds at 1 and 10 WAP (Supplemental Fig. S16). The 1 and 10 WAP stages correspond to the early stages of FA biosynthesis and the active biogenesis of LDs, respectively, as shown in Fig. 3, indicating that miRNAs may be involved in the regulation of genes related to these processes. Furthermore, among 123 oil-related genes identified in the transcriptome, 19 genes were targeted by at least one miRNA. These targets were scattered in different stages of oil accumulation (Supplemental Table S18).

CircRNAs have been reported to regulate gene expression in animals through different mechanisms, such as miRNA sponges, specific RNA-RNA interactions, and alternative splicing (Li et al., 2018). These genes were also found to influence the expression of their parental genes either positively(Tong et al., 2018, Wang et al., 2018) or negatively (Lu et al., 2015). In this study, 123 circRNAs were expressed throughout seed development, with higher read counts observed in 1 and 17 WAP (Supplemental Fig. S17A, B). Moreover, we found that the abundances of two circRNAs were positively correlated, whereas that of one circRNA was negatively correlated with the mRNA transcript abundances of their parental genes involved in oil biosynthesis (Supplemental Fig. S17C).

LncRNAs are longer than 200 nts and perform their biological functions through mechanisms such as epigenetic modification (Csorba et al., 2014, Heo and Sung, 2011), transcriptional regulation (Kim et al., 2010, Fatica and Bozzoni, 2014) and post-transcriptional regulation (Campalans et al., 2004, Dey et al., 2004). As shown in Supplemental Fig. S18, the expression of lncRNAs was stage-specific, with more genes expressed in the seeds at 1, 5 and 7 WAP, where most of the genes in the *Pull* and *Push* modules were expressed at high levels (Supplemental Fig. S18B; Fig. 3). Interestingly, even though fewer lncRNAs were expressed in the seeds at 10, 15 and 17 WAP, significantly more FPKMs of lncRNA were distributed in these stages (Supplemental Fig. S18A, C). Moreover, 19586 lncRNAs were expressed at least at one stage during seed development, and 70 of 123 oil-related genes were co-expressed with 1426 lncRNAs (Supplemental Table S19). Notably, while the vast majority of lncRNAs are expressed at markedly lower levels than mRNAs in our study (Supplemental Fig. S18C), several lncRNAs with co-expression patterns with core desaturase-encoding genes (*AAD6*, *FAD2*, and *FAD3*) exhibit high expressed at specific developmental stages (Fig. 5), which highlights their potential critical regulatory roles in the expression of these key desaturase genes.

**Fig. 5.**
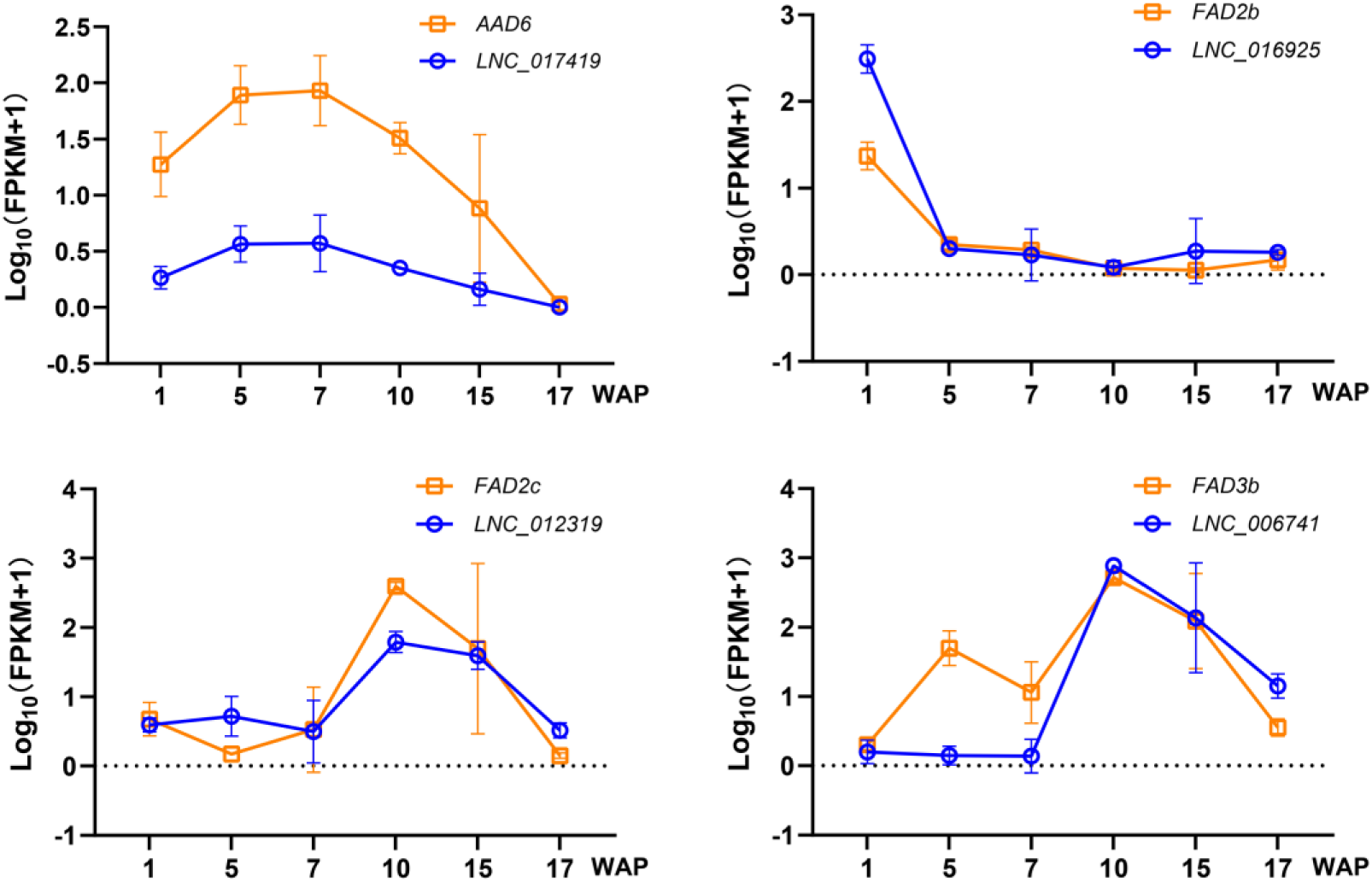
Expression profiles of genes encoding SAD, FAD2b, FAD2c, and FAD3b and their co-expressed lncRNAs.

## DISCUSSION

We report the first chromosome-scale reference genome for sacha inchi. Together with transcriptomic profiles from six key seed developmental stages, this resource systematically reveals the molecular basis for the high oil content and exceptionally high ALA content characteristic of its seeds. The underlying mechanism centers on the precise temporal coordination of the canonical “*Push*, *Pull*, *Package*, *Protect*” lipid assembly modules and multi-layered, specific regulation of the key desaturase gene.

### Conserved Lipid Synthesis Framework and Species-Specific Temporal Optimization

Seed oil synthesis in plants follows a highly conserved metabolic framework (Troncoso-Ponce et al., 2011), as depicted by Vanhercke et al. (2019). Our study demonstrated that sacha inchi has optimized this framework through a unique temporal strategy. Specifically, the *Push* module (de novo fatty acid synthesis) provides a continuous carbon flux throughout seed development, whereas the *Pull* module—particularly desaturation and esterification steps determining oil unsaturation—is specifically activated during the mid-to-late stages (10-15 weeks after pollination, WAP). This timing efficiently drives ALA synthesis and its incorporation into triacylglycerols (TAGs). Concurrently, the *Package* (lipid droplet formation) and *Protect* (lipolysis inhibition) modules are co-activated during the late accumulation phase, ensuring efficient storage and stability of the synthesized oil. This pattern of temporal regulation is also observed in tung tree (*Vernicia fordii*), which accumulates high levels of eleostearic acid (Zhang et al., 2019).

### The Central Regulatory Node for High ALA Accumulation: Transcriptional and Expression Control of FAD3

Our transcriptomic data clearly showed that the expression dynamics of *PvFAD3* are tightly synchronized with the ALA accumulation curve in seeds (Fig. 4), establishing transcriptional regulation as the core driver of this high-ALA trait. This regulatory logic is conserved across species: *AhFAD3-A01* expression levels in peanut directly determine seed ALA content in Arabidopsis and peanut (Li et al., 2025); in camelina, seed-specific expression of the exogenous *PfFAD3-1* gene elevates ALA content from ∼35% to 50% (Park et al., 2023). Functional validation further confirmed this finding: seed-specific overexpression of *PvFAD3* in tobacco significantly increased the proportion of ALA in seed oil (Liu et al., 2022, Yang et al., 2020). Collectively, these evidences establish *FAD3* as a key master gene for improving oil quality in oilseed crops.

Recent studies in other high-ALA crop species, such as tree peony, have provided deeper insight into the complex regulatory network governing *FAD3* and its pathway. In tree peony, *FAD3* expression is precisely controlled by a multi-layered, temporally defined transcriptional network. This includes global flux control driven by AP2 family transcription factors (e.g., *PrWRI1*) and a direct ‘switch’ mechanism for *FAD3* transcription, which is composed of multiple bZIP factors (e.g., the activator *PrABI5*, the activator *PsbZIP44*, and its repressor *PsbZIP9*) (Xie et al., 2023, Zhang et al., 2025, Xu et al., 2025). Furthermore, the transcription factor *PrDREB2D* ensures efficient channeling of ALA into storage oil by activating *PrPDAT2*, an acyltransferase with a substrate preference for ALA (Yang et al., 2024). Studies have reported that in sacha inchi seeds, the most highly unsaturated DAG(18:3/18:3) is preferentially used for TAG synthesis (Fu et al., 2024b).

While the active site ASP-104 in the PvFAD3 protein has been well-documented (Fu et al., 2025), our analysis of non-coding RNAs in sacha inchi reveals an additional layer of post-transcriptional regulation. For example, a specific lncRNA (e.g., *LNC_006741*) strongly co-expressed with *FAD2/FAD3*, whereas seed developmental stage-specific miRNAs and circRNAs may fine-tune early- and mid-phase genes in the lipid synthesis pathway through targeting or competitive endogenous RNA mechanisms. Together, these findings outline a multidimensional regulatory landscape of lipid synthesis compassing transcriptional activation, post-transcriptional fine-tuning by ncRNAs, and enzymatic modification.

## CONCLUSION

In conclusion, the high-quality, chromosome-scale genome assembly of sacha inchi presented in this study serves as a landmark resource for understanding the genetic architecture and evolutionary history of this unique woody oilseed crop. By integrating phylogenomic analysis with temporal transcriptomic profiling, we not only clarified ancient whole-genome duplication events within the Euphorbiaceae lineage but also elucidated the molecular mechanism governing the exceptional accumulation of ALAs in sacha inchi seeds. The identification of key desaturase genes, particularly the concerted action of *SAD*, *FAD2*, and *FAD3*, alongside the discovery of potential lncRNA-mediated regulatory layers, provides a comprehensive blueprint for future metabolic engineering. Furthermore, this genomic framework will be instrumental in accelerating molecular marker-assisted breeding and CRISPR-based genome editing to optimize oil profiles and enhance agronomic traits. Ultimately, our findings provide a foundation for the sustainable exploitation of sacha inchi, promoting it from a regional specialty to a globally significant source of high-value omega-3 fatty acids for the nutritional and pharmaceutical industries.

## MATERIALS AND METHODS

### Plant material

A one-year-old sacha inchi plant, which was grown at the Xishuangbanna Tropical Botanical Garden (21°54’N, 101°46’E, 580 m above sea level), Chinese Academy of Sciences, Mengla, Yunnan, was selected for whole genome sequencing. Genomic DNA was isolated from leaf tissue.

### Chromosome number of sacha inchi

To investigate the chromosome number of sacha inchi, fresh roots in a length of 1∼2 cm were harvested from plants that germinated from seeds. The roots were pretreated in 2 mM 8-hydrooxyquinoline at 20°C for 2 hours and then fixed with Carnoy’s solution (methanol: acetic acid = 3:1). Root tips were treated in a solution of 2% cellulase + 1% pectinase at 37°C for 60 minutes. Chromosome squashes were conducted on a precooled slide and stained with 4’-6-diamidino-2-phenylindole (DAPI) in an antifade solution (Vector). Images were captured using a Zeiss A2 fluorescence microscope (Carl Zeiss, Germany) with a micro CCD camera.

### Genome size estimation

Illumina reads (27.24 Gb of data, ∼39.34 X) from the 220 bp library were selected for the genome size estimation. This analysis was performed using “kmer_freq_stat” software developed by Biomarker Technologies. The formula used was genome size = k-mer_number/peak_depth. The genome size of sacha inchi was also estimated by flow cytometry using the genome of *Oryza sativa* L. ssp. *Japonica* (Goff et al., 2002) was used as the reference standard and propidium iodide (PI) was used as a fluorescent stain.

### Genome sequencing and assembly

An improved CTAB method (Murray and Thompson, 1980) was used to extract the genomic DNA. The modified CTAB extraction buffer included 0.1 M Tris-HCl, 0.02 M EDTA, 1.4 M NaCl, 3% (w/v) CTAB and 5% (w/v) PVP K40. Beta-mercaptoethanol was added to the CTAB extraction buffer to ensure DNA integrity and quality. We sequenced 15 different libraries with different insert sizes, including two paired-end libraries with the short insert sizes of 220 bp and 500 bp, and twelve mate-pair libraries with insert sizes of 3 kb, 4 kb, 8 kb, 10 kb, 15 kb, and 17 kb (Supplemental Table S1). PacBio sequencing was performed on a PacBio RS-II platform with C4 chemistry to obtain long reads.

The sacha inchi genome assembly was first performed using Illumina reads with ALLPATHS-LG software (Gnerre et al., 2011) with the default parameters, contigs were combined into scaffolds with SSPACE v2.3 (Boetzer et al., 2011), and gaps within the scaffolds were filled with GapCloser v1.12 (Luo et al., 2012) in the SOAPdenovo package (Luo et al., 2012). Based on the preassembly, we utilized PBJelly v14.9.9 (English et al., 2012) to perform gap filling with the corrected PacBio reads

### Hi-C assisted genome assembly

To get a high-resolution genome contact map, we used *in situ* Hi-C following the protocol of a previous study (Rao et al., 2014) with some modifications. The fresh plant leaves were crosslinked with 1% formaldehyde. To destroy the cell wall, formaldehyde fixed powder was added to buffer solution. The restriction endonuclease MboI was used to digest DNA, followed by biotinylated residue labeling. The Hi-C library was then sequenced on BGISEQ-500 platform with 100 bp pair-end sequencing. HiC-Pro pipeline (Servant et al., 2015) was implemented in quality control. Of all 864,873,129 raw pair-end reads, there are 17% (147,193,076) paired Hi-C reads are valid and suitable for following analysis (Supplemental Fig. S3). Basing on these valid Hi-C reads, we used Juicer (Durand et al., 2016) and Aiden’s Hi-C assembly pipeline (Dudchenko et al., 2017) to assemble the genome with the main parameter “-m haploid -s 4 -c 29”. As tiny scaffolds (shorter than 500 bp) in original genome have fewer Hi-C contacts and are difficult to analyze, we removed these scaffolds from our final assembly.

### Genome annotation

The repeat sequences were annotated via various methods. Tandem repeats were searched across the genome using the software Tandem Repeats Finder (4.07) (Benson, 1999). Transposable elements (TEs) were predicted using a combination of homology-based comparisons with RepeatMasker (4.0.6) and protein-based RepeatMasking (http://www.repeatmasker.org), and via *de novo* approaches with LTR_FINDER (1.0.6) (Xu and Wang, 2007) and RepeatModeler (1.0.8) (http://www.repeatmasker.org). *De novo* generated TEs were also classified into TE families by RepeatMasker.

Protein coding genes were predicted based on a combination of ab initio, conserved protein homologs and assembled transcripts. AUGUSTUS (3.1) (Stanke et al., 2008) and GlimmerHMM (3.0.4) (Majoros et al., 2004) were used for ab initio prediction. AUGUSTUS was trained with proteins of *Hevea brasiliensis* and GlimmerHMM was run with pre-trained parameters specialized for *A. thaliana*. We predicated genes from repeat-masked genome sequences. Protein sequences of *A. thaliana* [The *A. thaliana* Information Resource (TAIR), www.arabidopsis.org], *H. brasiliensis* (Tang et al., 2016)*, J. curcas* [were assembled by using the Hi-C sequencing data we generated and raw data from (Wu et al., 2015)]*, L. usitatissimum* (Phytozome v12.1, https://phytozome.jgi.doe.gov/pz/portal.html)*, Manihot esculenta* (Bredeson et al., 2016)*, Populus trichocarpa* (Tuskan et al., 2006)*, R. communis* (Chan et al., 2010)*, V. fordii* (Cui et al., 2018) and *Vitis vinifera* (Jaillon et al., 2007) were compared against sacha inchi genome using BLAT, and the best hits were used as input of GeneWise (2.4.1) (Birney et al., 2004) to predict gene models. Transcripts were assembled using Trinity (2.6.6) (Grabherr et al., 2011). All the results from the three types of predictions were integrated by GLEAN (Elsik et al., 2007).

Non-coding RNAs were identified by searching against various RNA libraries. tRNAscan-SE (1.3.1) (Lowe and Eddy, 1997) was run on the assembled genomic sequence to identify tRNAs. rRNA sequences were annotated based on homology to previously published rRNA sequences in plants. snRNAs and miRNAs were annotated by employing the INFERNAL (1.1.1) software to search against the Rfam (version 12.0) database (Kalvari et al., 2018). “cmsearch” programs in INFERNAL were used to identify the non-coding RNAs with an e-value cutoff of 0.01 (Kalvari et al., 2018).

We chose 1:1 single-copy families to construct the phylogenetic tree. Proteins and CDSs from single-copy families of 13 selected species were aligned using MUSCLE (3.8.31) (Edgar, 2004b). Four-fold degenerate sites were extracted and merged into a supergene as an input of MrBayes (3.1.2) (Huelsenbeck and Ronquist, 2001) with *O. sativa* as the outgroup. The divergence time was estimated by MCMCTree from the PAML (4.4) (Yang, 2007) package based on HKY85 model. Correlated rates were used for molecular clock model. Seven calibration fossil evidences were from http://www.timetree.org.

The expansion and contraction of gene families were determined by comparing the family size differences between the most recent common ancestor (MRCA) and each of the 13 plant genomes using the CAFÉ (2.1) program (De Bie et al., 2006). A random birth and death model was used to study changes in gene family size along each branch of the phylogenetic tree. Using conditional likelihoods as the test statistics, we calculated the corresponding *p* values in each branch. A *P* value of 0.05 was used to identify families that were significantly changed in sacha inchi genome. Genes of expanded families were used for KEGG enrichment analysis.

For whole genome duplication and microsynteny, paralogs of each species were identified by an all-versus-all BLASTP approach with an e-value cutoff of 1e-5. Second, syntenic blocks were identified by MCscan (0.8) (Tang et al., 2008) based on paralogs (e-value < 1e-5 and number of genes required to call synteny >= 5). For each paralogous gene in syntenic blocks, Ks values were calculated using KaKs_calculator (2.0) on the basis of the CDS and protein alignment (Wang et al., 2010), and corrected fourfold degenerate transversion (4DTv) rates were calculated by self-determination program. Ks and 4DTv distributions were compared among six Euphorbiaceae species and *V. vinifera*.

### Transcriptome sequencing and analysis of seeds in six development stages

Seeds at 1, 5, 7, 10, 15 and 17 WAP (weeks after pollination) were collected for RNA extraction. After library preparation, they were sequenced on an Illumina HiSeq 2500 platform, and 125 bp paired-end reads were generated. Paired-end clean reads were aligned to the reference genome using HISAT2 (Langmead and Salzberg, 2012) v2.0.4. The mapped reads of each sample were assembled by StringTie (v1.3.3) (Langmead and Salzberg, 2012) in a reference-based approach. Gene expression was measured using the fragments per kilobase of transcript sequence per millions of base pairs sequenced (FPKM) method (Trapnell et al., 2014).

### Genes involved in FA and TAG biosynthesis

Genes involved in FA and TAG biosynthesis in *A. thaliana* (Li-Beisson et al., 2013) were retrieved from *A. thaliana* proteome and searched against sacha inchi proteomes, using BLASTP with an E-value of 1e-10. The resulting hits were then mapped back to the *A. thaliana* proteome, and the most similar protein was selected according to the *P*-value, following suggested steps in (Samach, 2013).

## ACKNOWLEDGMENT

We thank the Central Laboratory of the Chinese Academy of Sciences Xishuangbanna Tropical Botanical Garden for providing the research facilities. This work was supported by the Youth Talent Support Program of Yunnan Province (YNWR-QNBJ-2020-172), the West Light Foundation of the Chinese Academy of Sciences, the Guangxi Specific Project for Science and Technology Bases and Talents (AD23026337), the National Natural Science Foundation of China (31870291), the Natural Science Foundation of Yunnan Province (2016FB051), and the CAS 135 program (2017XTBG-T02).

## AUTHOR CONTRIBUTIONS

BZP, ZFX, CL and TL conceived the project. QF, MSC, YBT and HH prepared the sacha inchi plants, designed the sample collection methodology and participated in manuscript revisions. XZ, XDH and CL led the genome analysis, conducted the genome assembly, and predicted the gene structure and repeat sequences. BZP, XDH and LJN conducted the flow cytometry analysis. YS and ZC investigated the chromosome number of sacha inchi. All of the authors listed above participated in discussions about the project and data. BZP and ZFX wrote the manuscript, and all the authors helped with manuscript revisions. All the authors read and approved the final manuscript.

## CONFLICT OF INTEREST

The authors declare that they have no competing interests.

## DATA AVAILABILITY STATEMENT

The sacha inchi genome and RNA-seq sequences have been deposited in GenBank under the accession numbers of JBLVOQ000000000 and PRJNA1175046, respectively.

## DECLARATIONS WITH ETHICS STATEMENT

The *Plukenetia volubilis* used in this study is not an endangered or protected species, and no specific permissions were required for the study.

## SUPPORTING INFORMATION

Additional Supporting Information may be found in the online version of this article.

**Fig. S1** Overview of sacha inchi plant. (A) Sacha inchi plant. (B) Fruits. (C) Seeds. (D) Raw seed oil.

**Fig. S2** K-mer analysis of the sacha inchi genome. The figure shows the frequency of 19 k-mers which are 19 bp sequences from the reads (after filtering) of short-insert size libraries.

**Fig. S3** Statistics of read pairs obtained through Hi-C sequencing alignment of restriction fragments.

**Fig. S4** Flow cytometry estimation of the sacha inchi genome size compared with the reference standard genome size of *O. sativa* L. ssp. *Japonica*.

**Fig. S5** GC content and sequencing-depth analyses. The abscissa represents the GC content, and ordinate represents the average depth.

**Fig. S6** Insertion dates of LTR transposons in the genomes of sacha inchi*, H. brasiliensis* and *M. esculenta*.

**Fig. S7** Seeds at different development stages selected for transcriptome sequencing. WAP, weeks after pollination.

**Fig. S8** Comparisons of gene length, CDS length, exon length, intron length and exon number in sacha inchi and selected species.

**Fig. S9** Orthology delineation of the protein-coding gene family repertoires of 13 selected species.

**Fig. S10** Expression profiles of genes encoding lipoxygenases (LOXs) and 12-oxophytodienoate reductases (OPRs), which are enriched in the oxidative branch leading to jasmonic acid (JA) biosynthesis. The color scale shows the log10 (RPKM+1) values.

**Fig. S11** Synteny between sacha inchi and *J. curcas.* Syntenic blocks were identified by MCscan. The coordinates of these syntenic blocks were plotted to display the microsynteny of the two genomes.

**Fig. S12** Synteny between sacha inchi and *M. esculenta.* Syntenic blocks were identified by MCscan. The coordinates of these syntenic blocks were plotted to display the microsynteny of the two genomes.

**Fig. S13** Profiles of the fatty acid composition of sacha inchi seed oil during seed development (Wang et al., 2014). PL, phospholipid; FFA, free fatty acid; NL, neutral lipid; C16:0, palmitic acid; C18:0, stearic acid; C18:1, oleic acid; C18:2, linoleic acid; C18:3, linolenic acid.

**Fig. S14** Expression profiles of genes encoding TAG lipases. The full names of the genes corresponding to the abbreviations are listed in Supplemental Table S16. The color scale shows log_10_ (RPKM+1) values.

**Fig. S15** Expression profiles of genes involved in the ALA metabolism pathway. The full names of the genes corresponding to the abbreviations are listed in Supplemental Table S16. The color scale shows log_10_ (RPKM+1) values.

**Fig. S16** Summary of miRNAs identified during seed development. (A) Heatmap of miRNAs. The color scale shows the log_10_ (TPM+1) values. (B) The miRNA number and total read counts of each stage.

**Fig. S17** Summary of ceRNAs identified during seed development. (A) Heatmap of ceRNAs. The color scale represents log_10_ (TPM+1) values. (B) The ceRNA number and total read counts of each stage. (B) The expression profiles of three ceRNAs and their parental genes.

**Fig. S18** Summary of lncRNAs identified during seed development. (A) Heatmap of lncRNAs. The color scale represents log_10_ (RPKM+1) values. (B-C) The number (B) and FPKM distribution (C) of mRNAs and lncRNAs at each stage.

**Table S1** Sequencing libraries and data yields for Illumina sequencing.

**Table S2** Summary of clean reads after filtering the raw reads from PacBio sequencing.

**Table S3** Length distribution of PacBio sequencing subreads.

**Table S4** Summary of sequences anchored to chromosomes.

**Table S5** Statistics of genome coverage.

**Table S6** Evaluation of the genome completeness of sacha inchi*, J. curcas* and *R. communis*.

**Table S7** Statistics of the repeat sequences.

**Table S8** Classification of the repeat sequences based on different methods.

**Table S9** Summary of protein-coding gene prediction.

**Table S10** Protein-coding gene and genomic GC content comparisons among different plant species.

**Table S11** BUSCO assessment of gene prediction.

**Table S12** Statistics for functional annotations.

**Table S13** Prediction of non-coding RNAs in the sacha inchi genome.

**Table S14** Statistics of gene family clusters.

**Table S15** KEGG enrichment analysis of genes within the significantly expanded families.

**Table S16** Expression profiles of genes involved in lipid biosynthesis pathways.

**Table S17** Data details of Fig. 4.

**Table S18** Differentially expressed miRNAs and their targeted oil-related genes.

**Table S19** Genes involved in lipid biosynthesis pathways and their co-expressed lncRNAs.

## Notes

### Competing Interest Statement

The authors have declared no competing interest.

