## Supplemental Figures for "Chromosome-scale genome of the woody oilseed crop sacha inchi elucidates the molecular basis of alpha-linolenic acid biosynthesis and triacylglycerol accumulation in seeds"

**Fig. S1** **Overview of sacha inchi plant**

**(A)** Sacha inchi plant. **(B)** Fruits. **(C)** Seeds. **(D)** Raw seed oil.

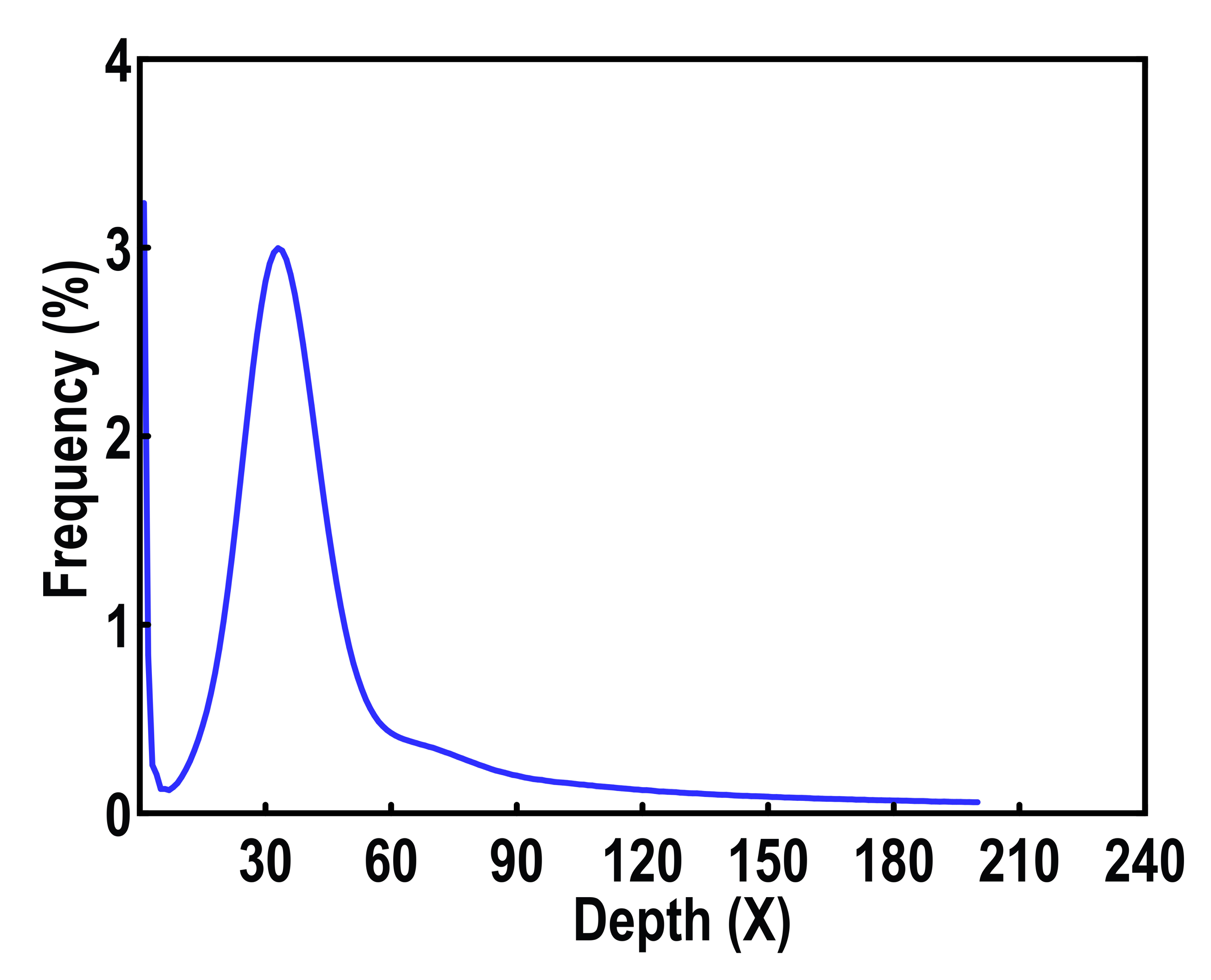

**Fig. S2** K-mer analysis of the sacha inchi genome

The figure shows the frequency of 19 k-mers which are 19 bp sequences from the reads (after filtering) of short-insert size libraries.

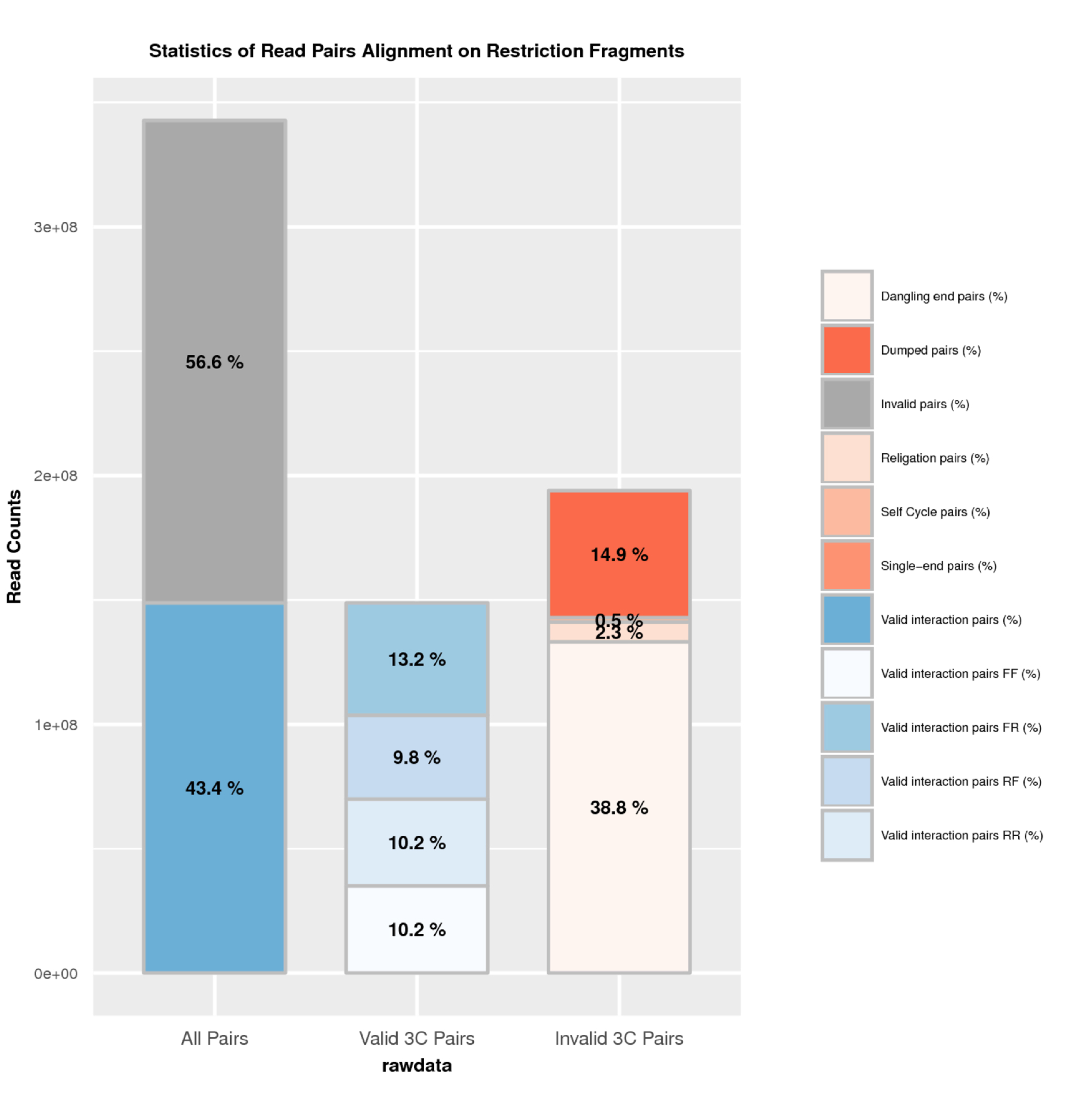

**Fig. S3** Statistics of read pairs obtained through Hi-C sequencing alignment of restriction fragments

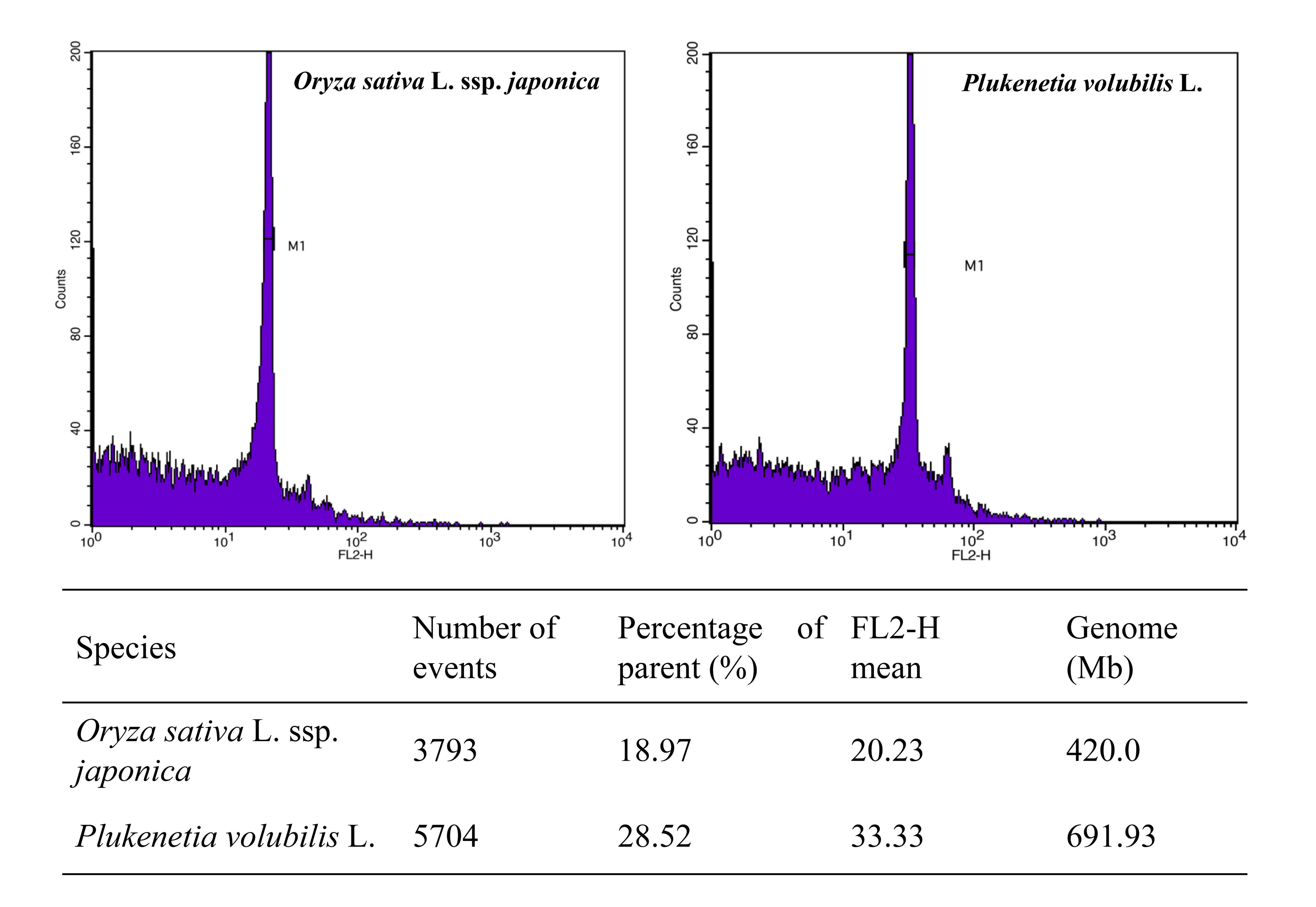

**Fig. S4** Flow cytometry estimation of the sacha inchi genome size compared with the reference standard genome size of *O. sativa* L. ssp. *Japonica*

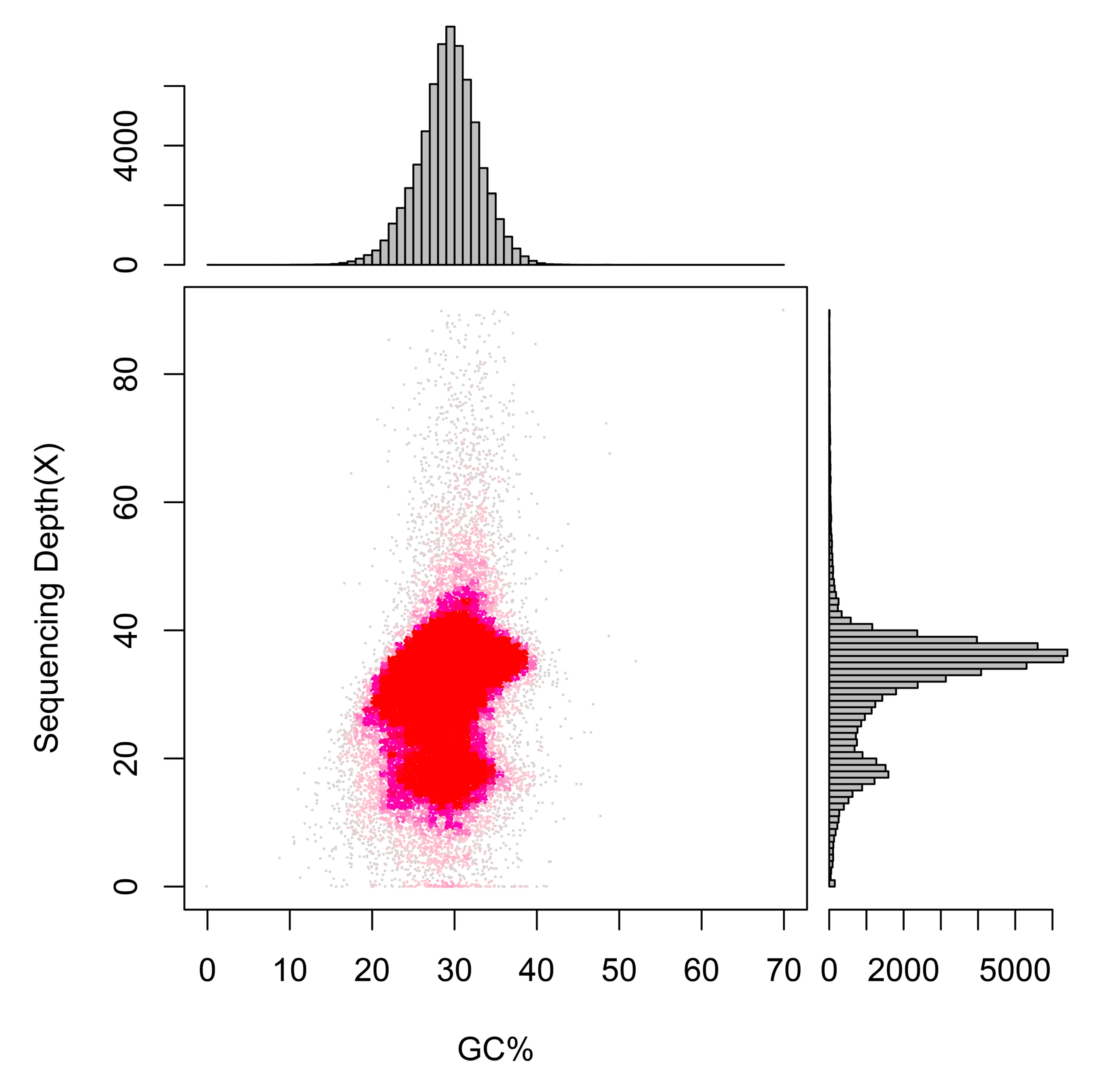

**Fig. S5** GC content and sequencing-depth analyses

The abscissa represents the GC content, and ordinate represents the average depth.

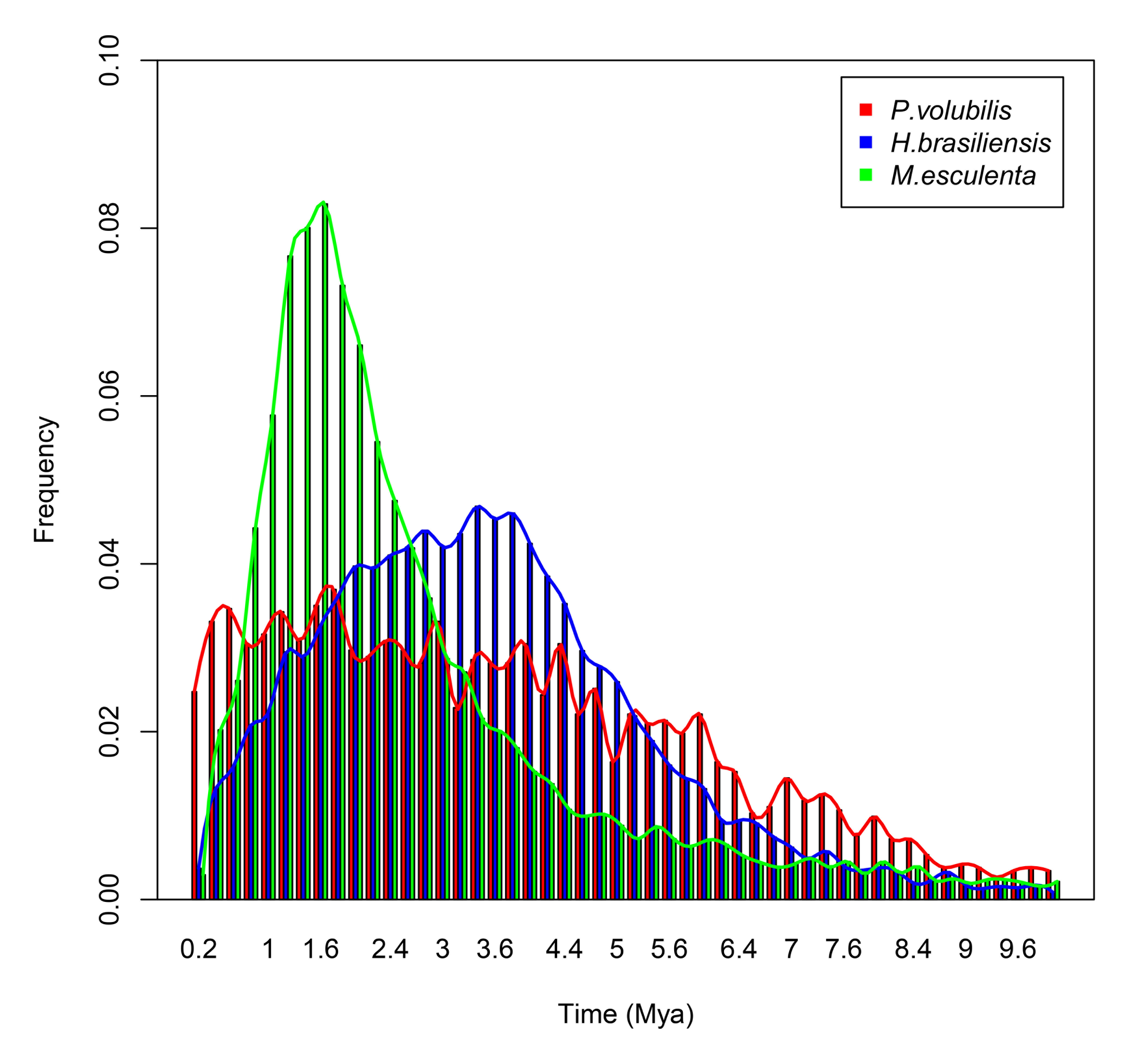

**Fig. S6** Insertion dates of LTR transposons in the genomes of sacha inchi*,* *H. brasiliensis* and *M. esculenta.*

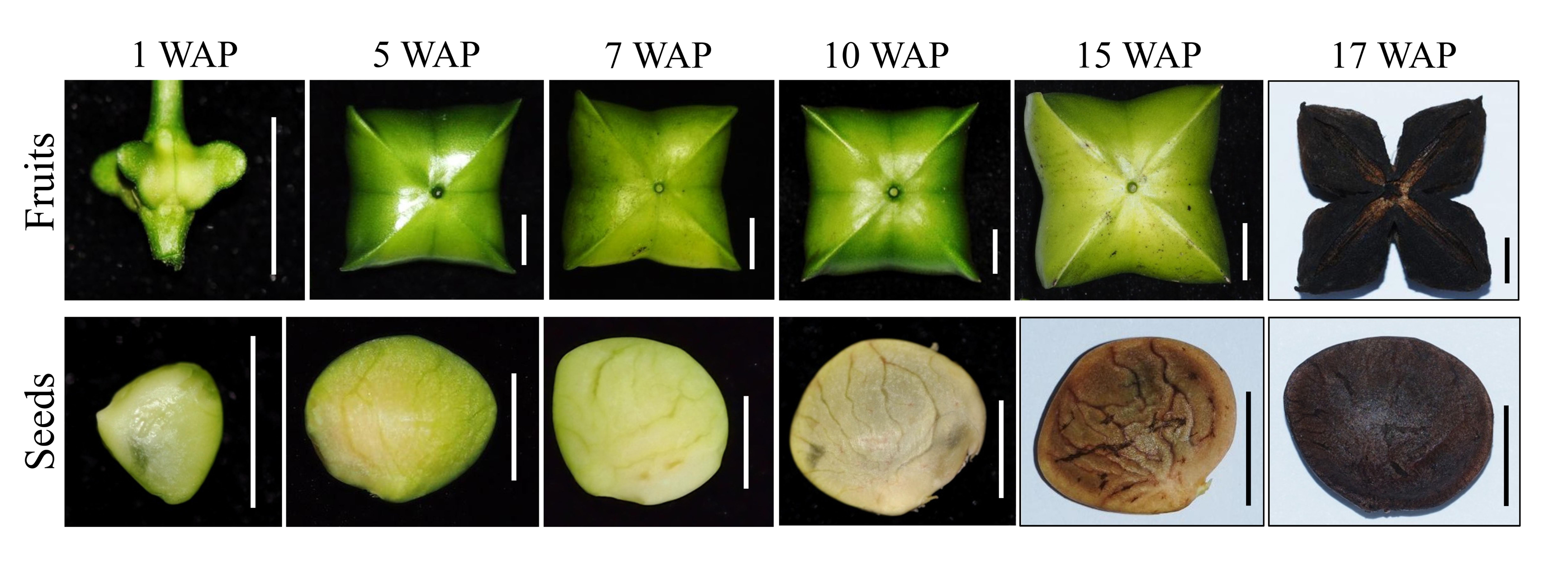

**Fig. S7** Seeds at different development stages selected for transcriptome sequencing WAP, weeks after pollination.

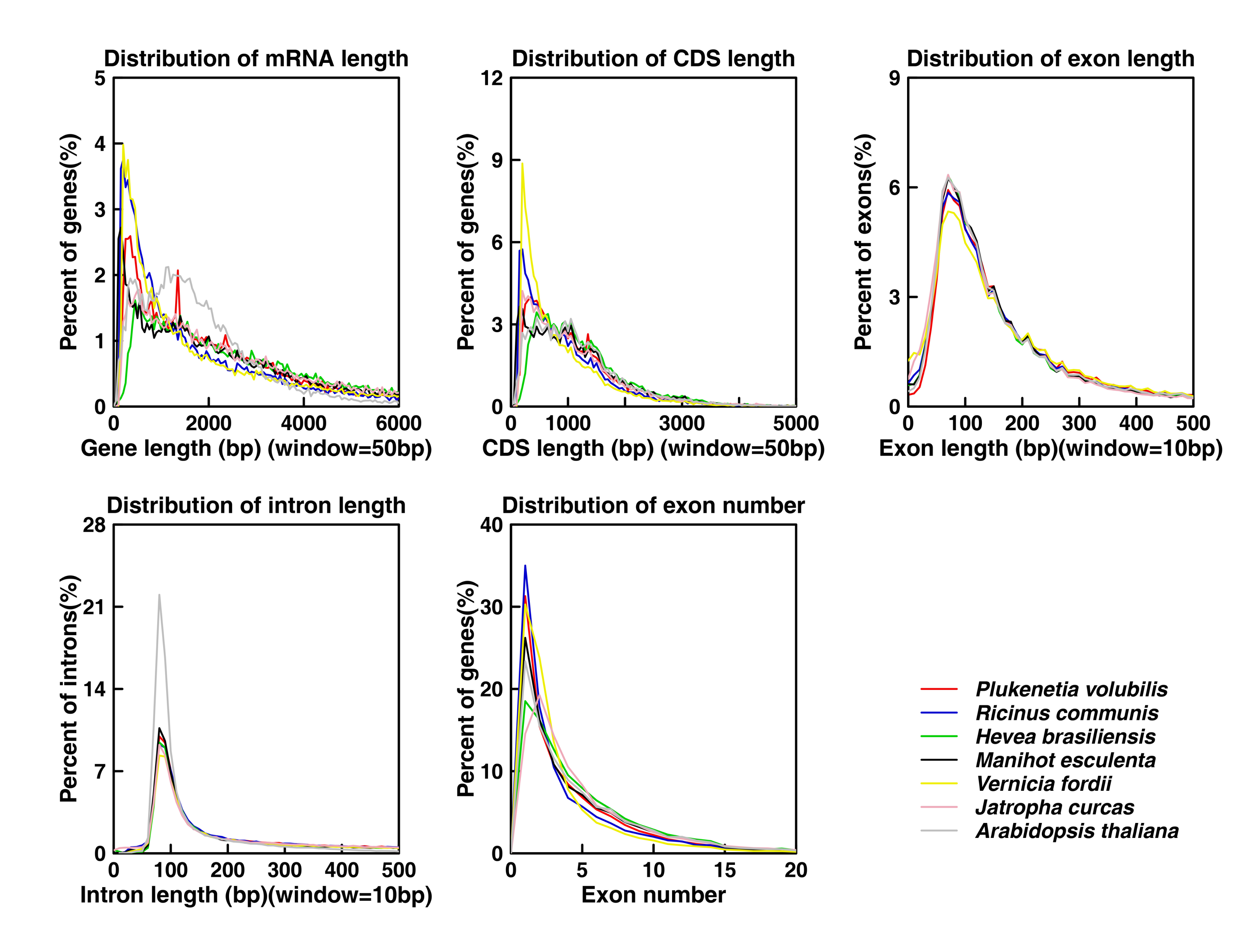

**Fig. S8** Comparisons of gene length, CDS length, exon length, intron length and exon number in sacha inchi and selected species.

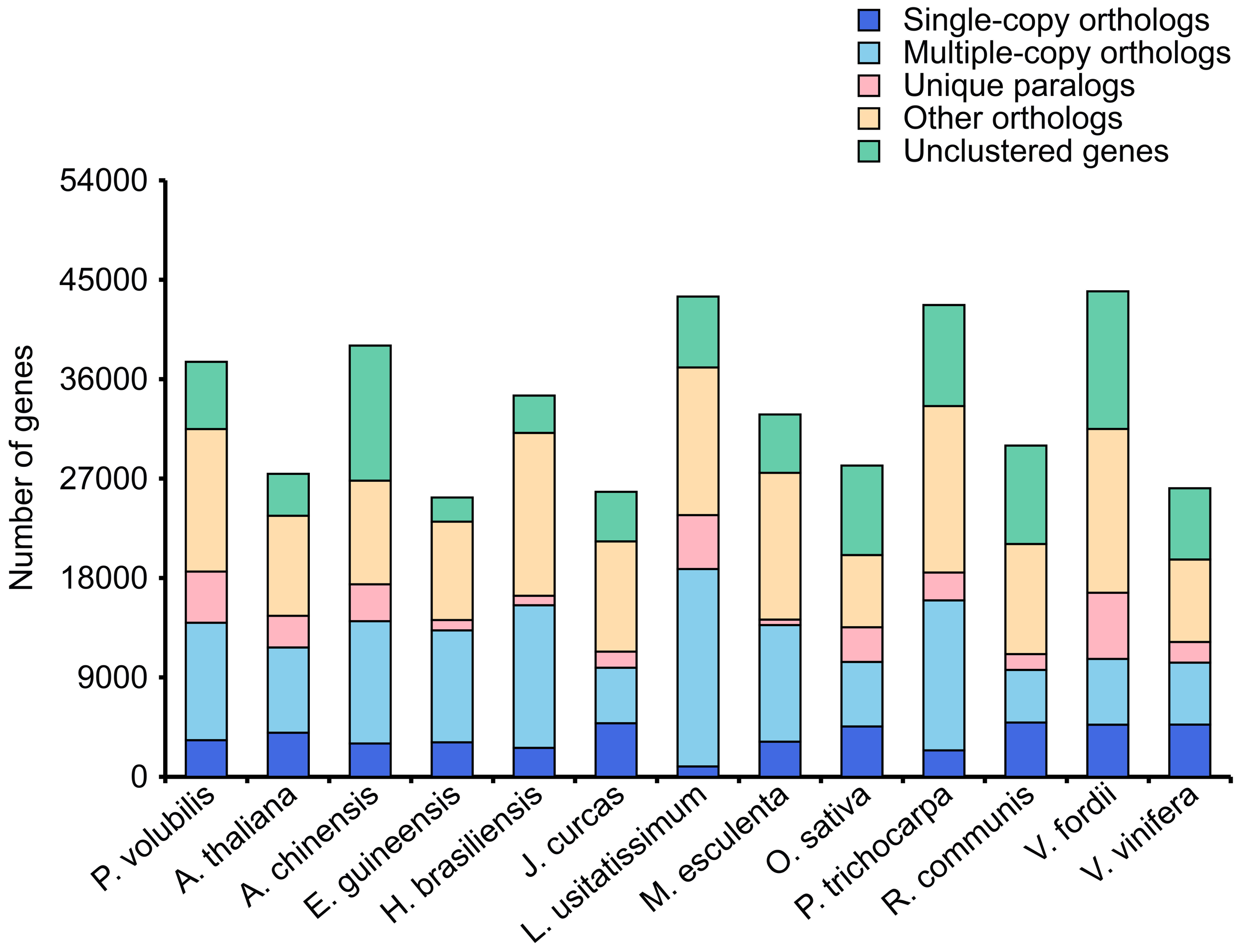

**Fig. S9** Orthology delineation of the protein-coding gene family repertoires of 13 selected species

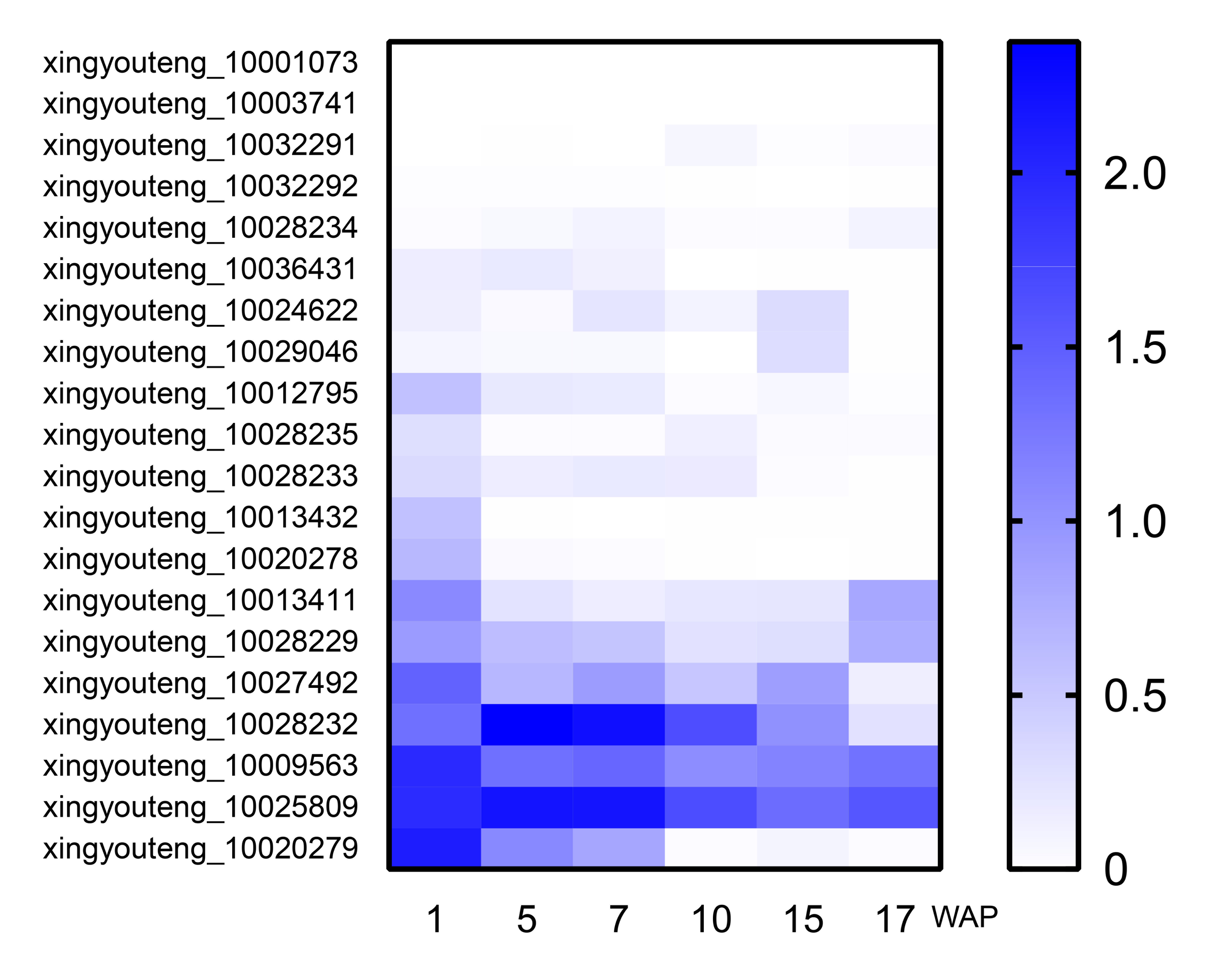

**Fig. S10** Expression profiles of genes encoding lipoxygenases (LOXs) and 12-oxophytodienoate reductases (OPRs), which are enriched in the oxidative branch leading to jasmonic acid (JA) biosynthesis. The color scale shows the log10 (RPKM+1) values.

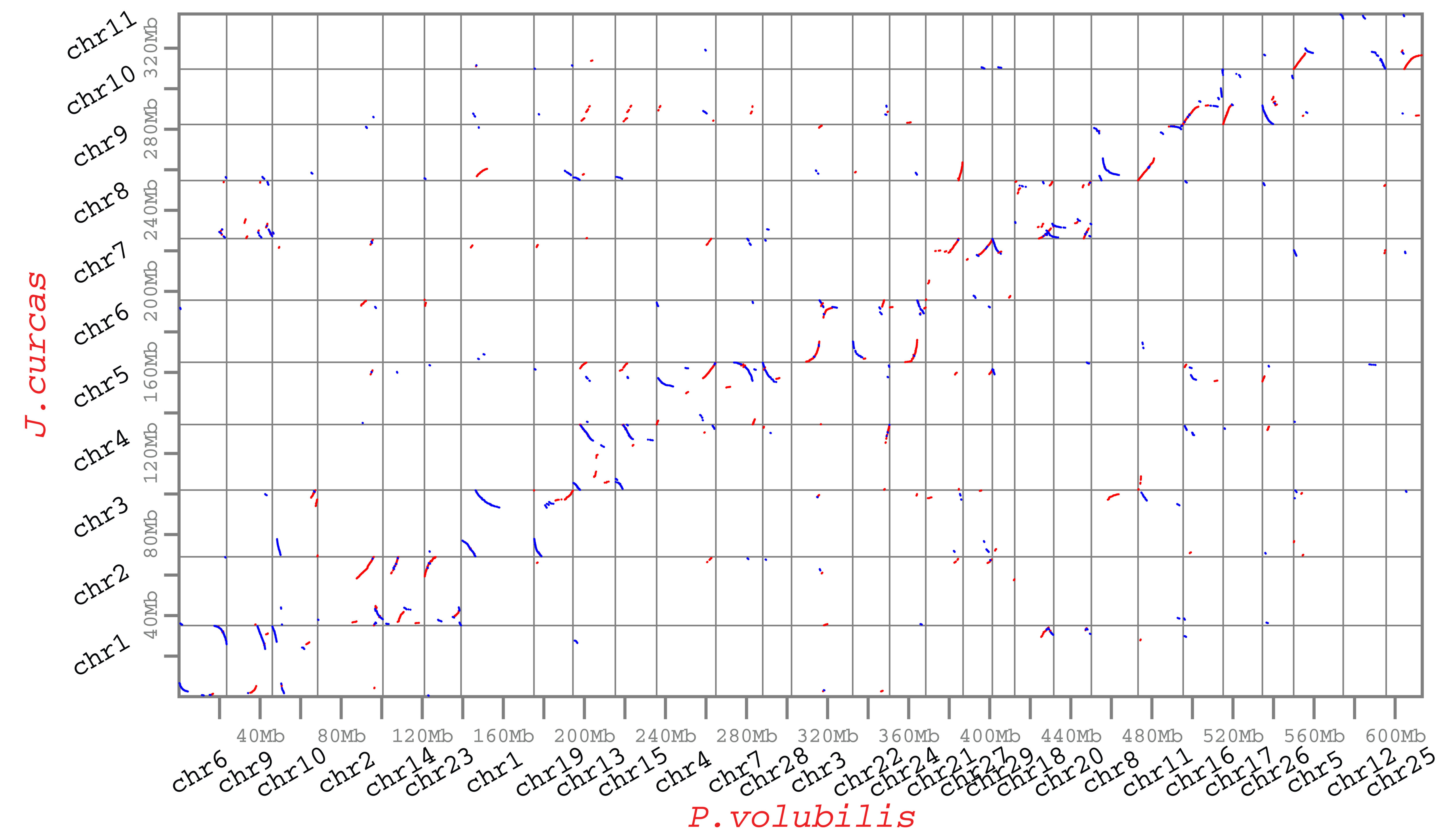

**Fig. S11** Synteny between sacha inchi and *J. curcas.* Syntenic blocks were identified by MCscan

The coordinates of these syntenic blocks were plotted to display the microsynteny of the two genomes.

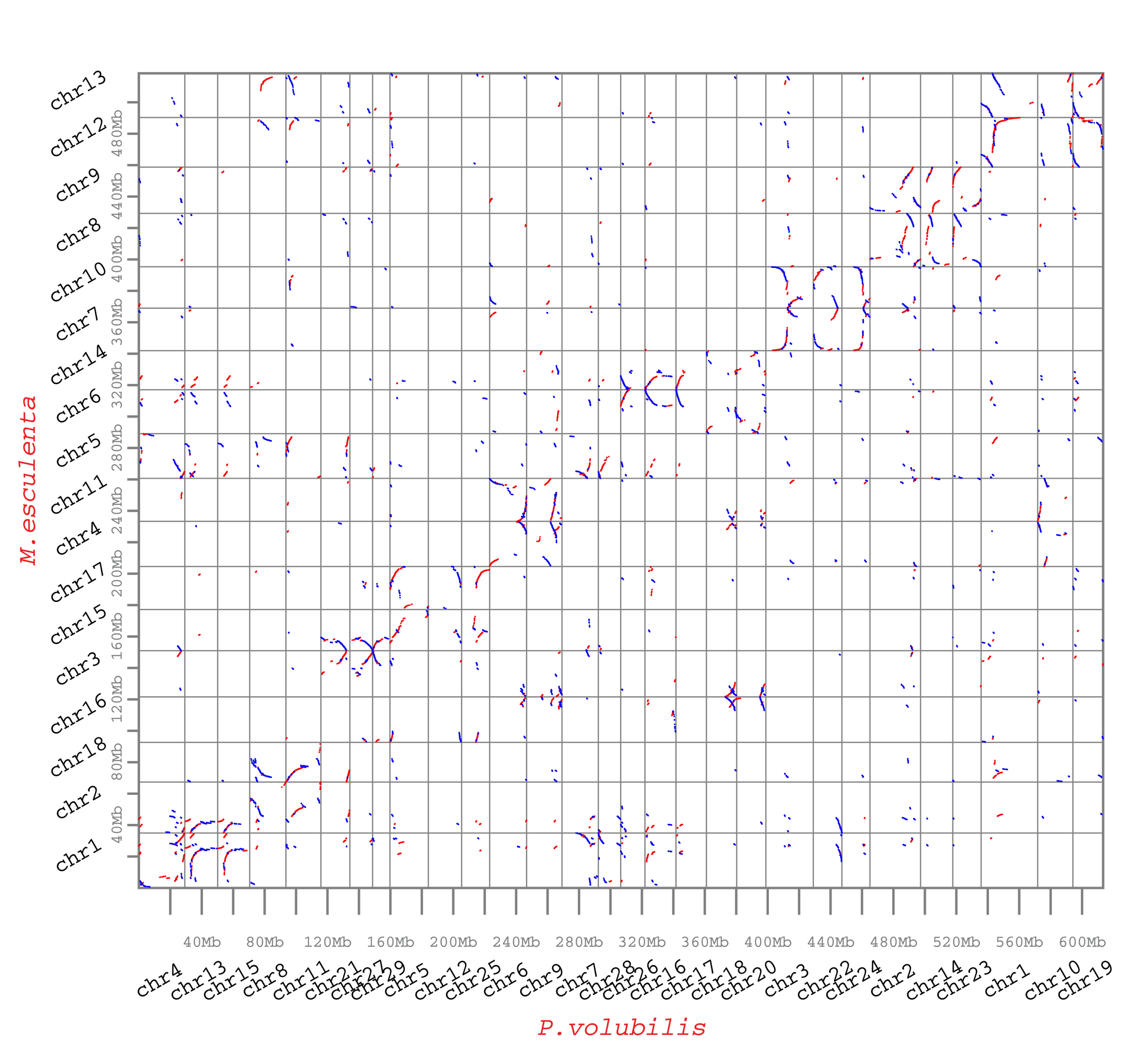

**Fig. S12** Synteny between sacha inchi and *M. esculenta.* Syntenic blocks were identified by MCscan

The coordinates of these syntenic blocks were plotted to display the microsynteny of the two genomes.

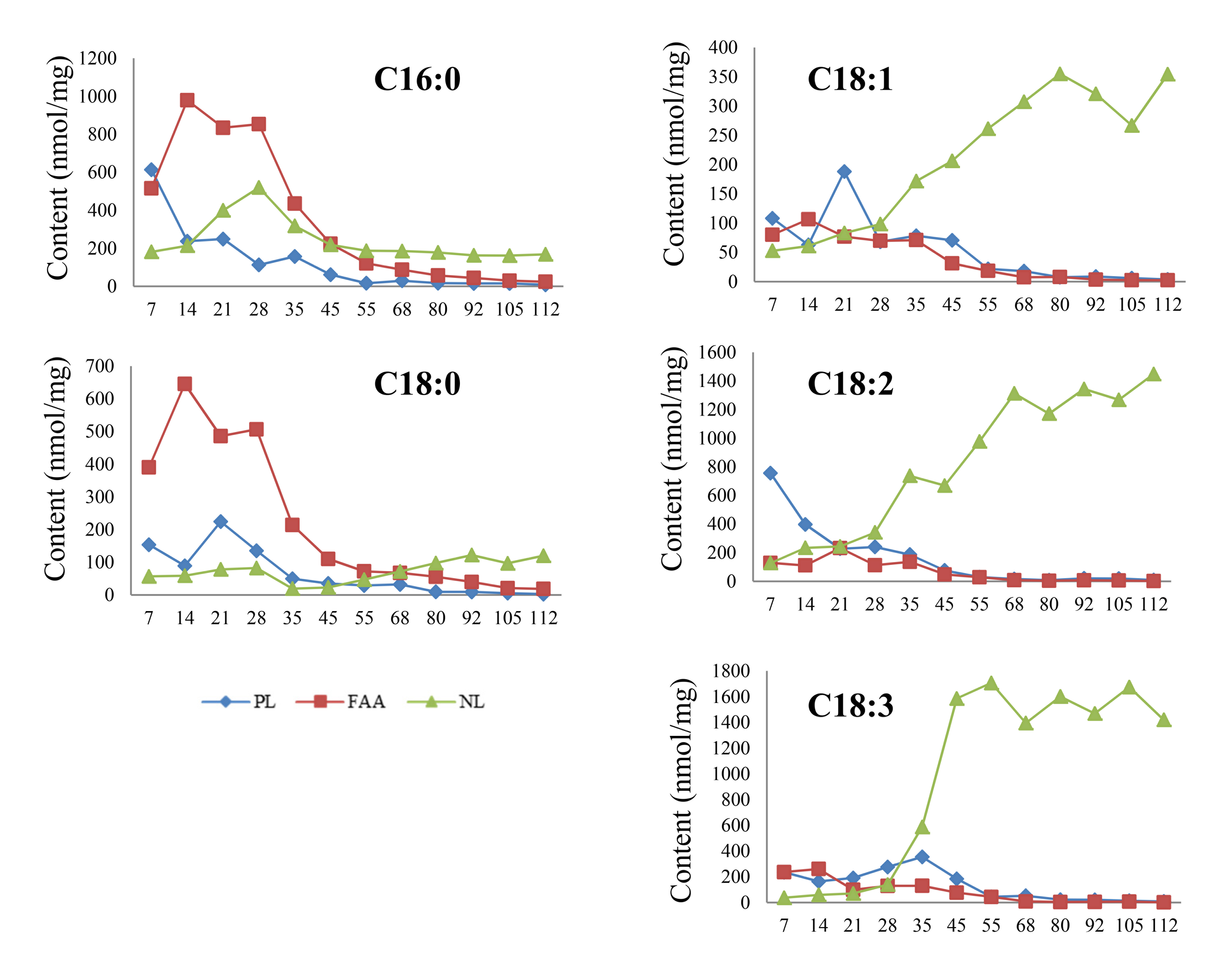

**Fig. S13** Profiles of the fatty acid composition of sacha inchi seed oil during seed development (Wang et al., 2014)

PL, phospholipid; FFA, free fatty acid; NL, neutral lipid; C16:0, palmitic acid; C18:0, stearic acid; C18:1, oleic acid; C18:2, linoleic acid; C18:3, linolenic acid.

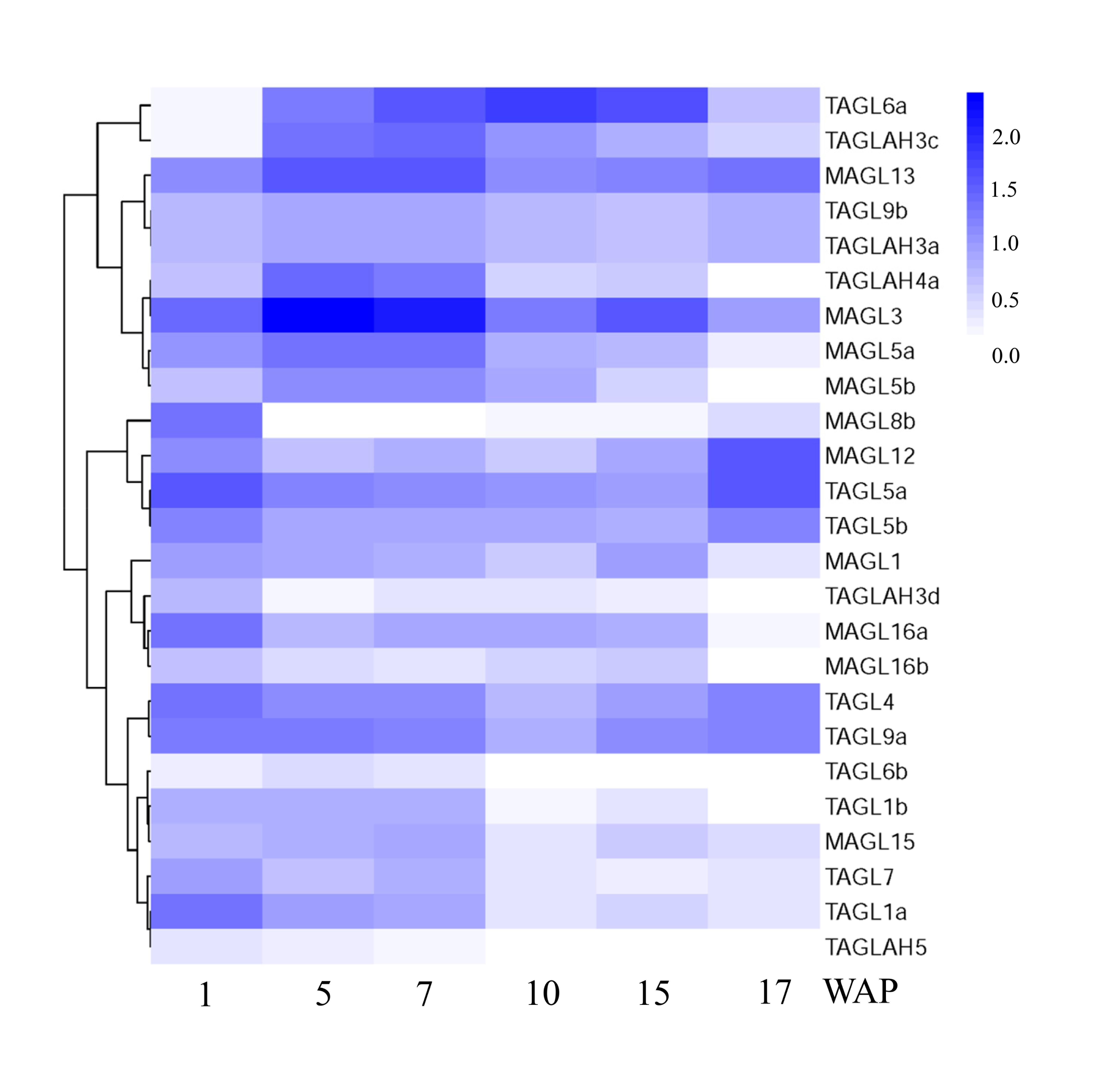

**Fig. S14** Expression profiles of genes encoding TAG lipases

The full names of the genes corresponding to the abbreviations are listed in Supplemental Table 16. The color scale shows log_10_ (RPKM+1) values.

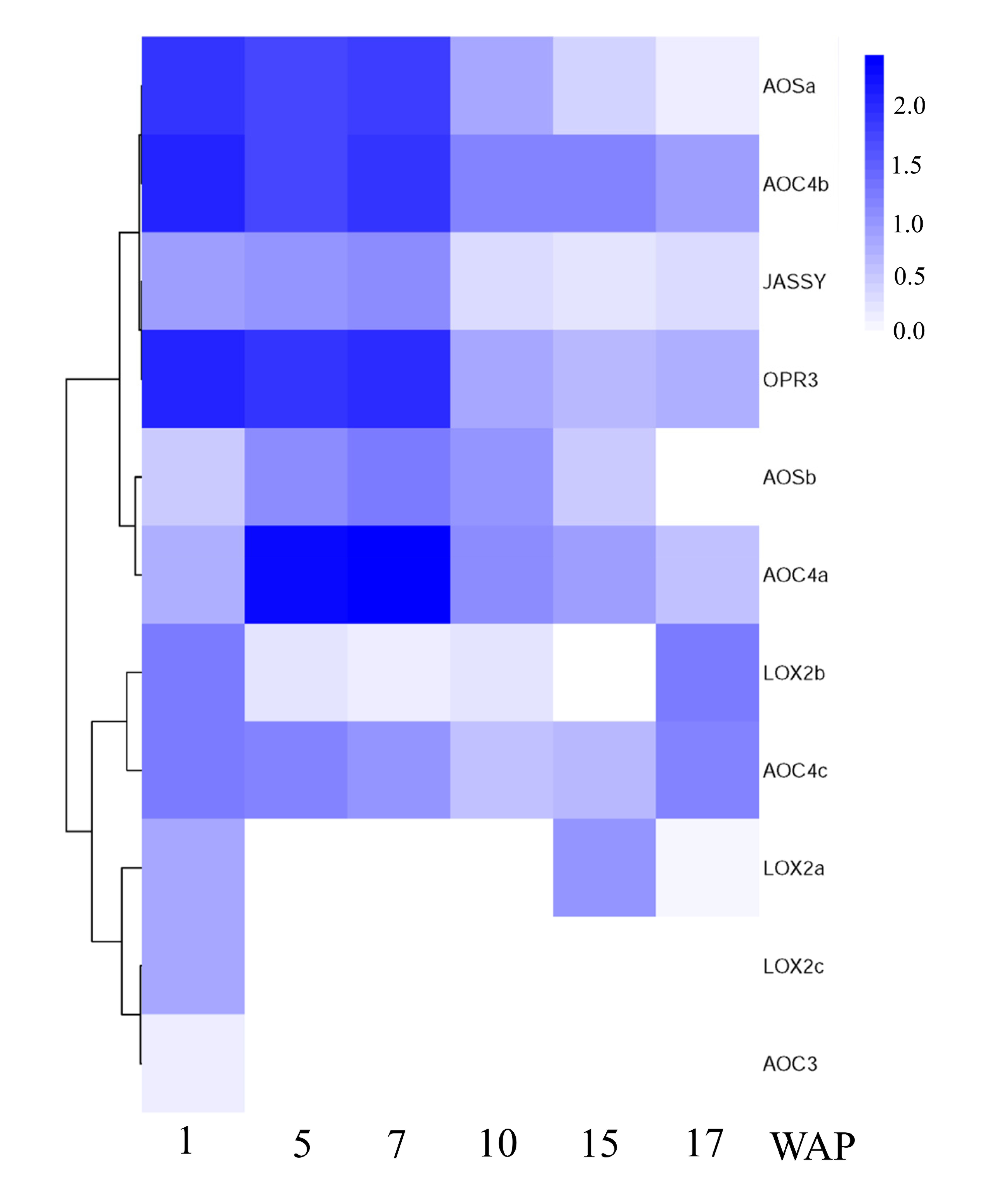

**Fig. S15** Expression profiles of genes involved in the ALA metabolism pathway

The full names of the genes corresponding to the abbreviations are listed in Supplemental Table 16. The color scale shows log_10_ (RPKM+1) values.

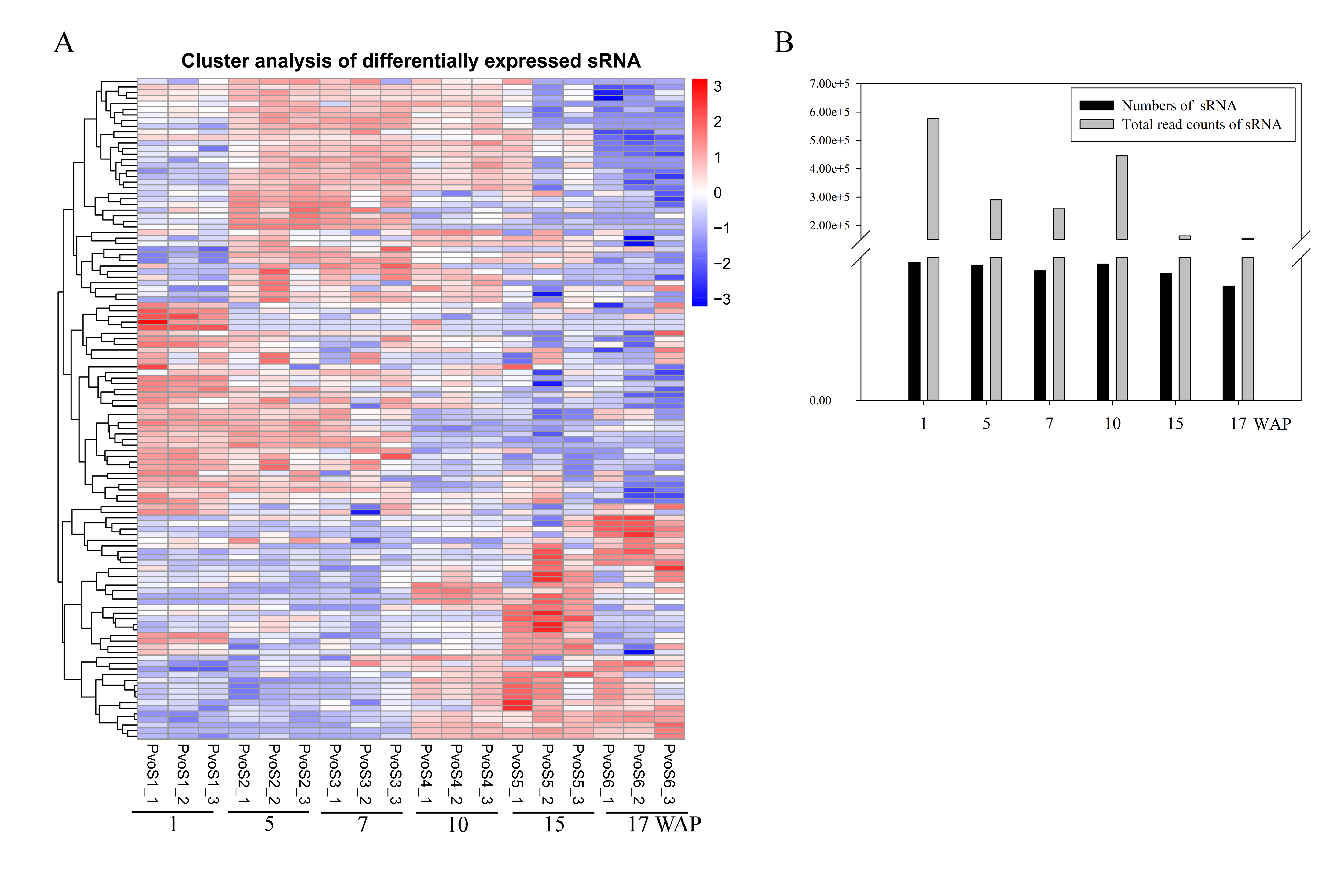

**Fig. S16** Summary of miRNAs identified during seed development

(A) Heatmap of miRNAs. The color scale shows the log_10_ (TPM+1) values. (B) The miRNA number and total read counts of each stage.

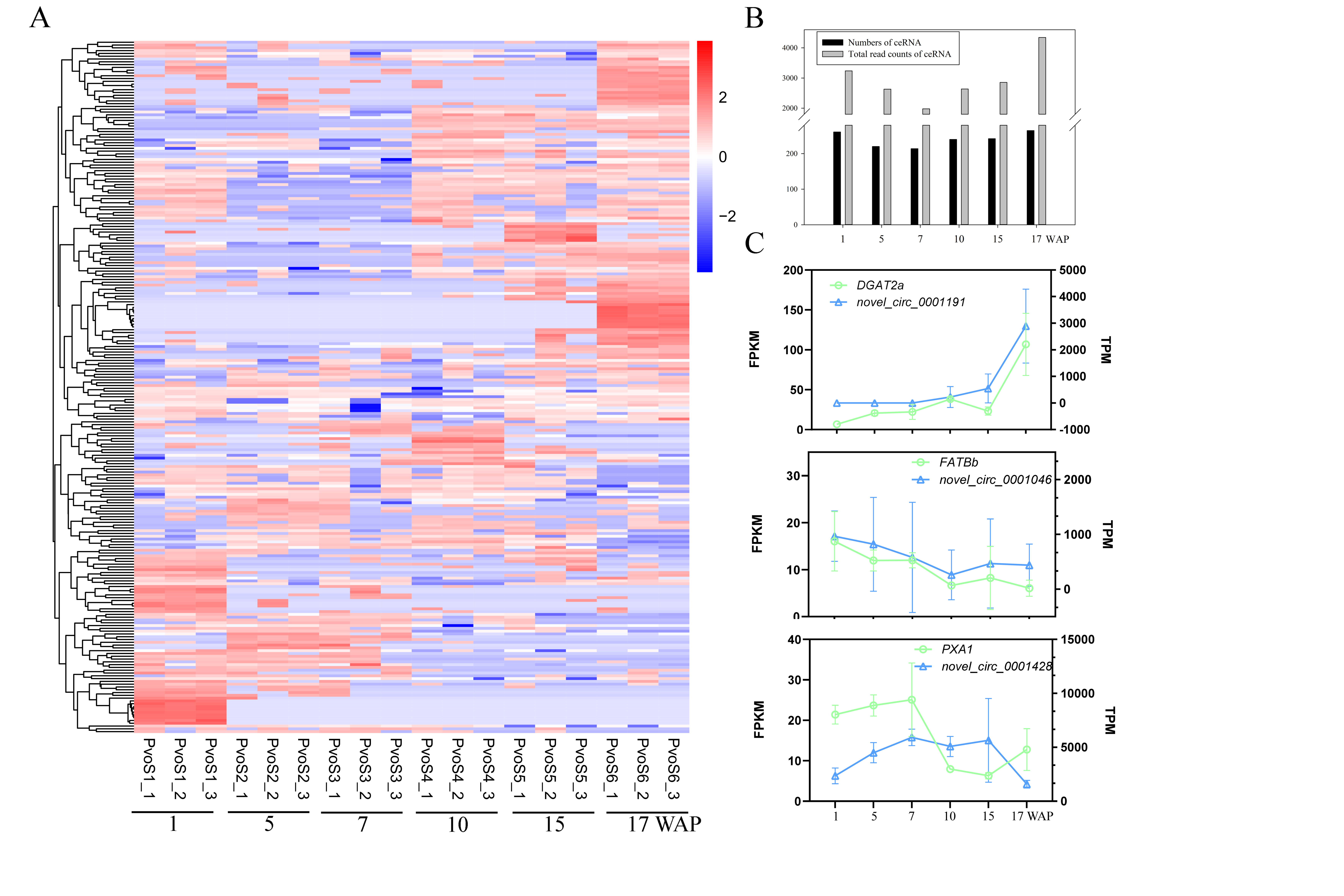

**Fig. S17** Summary of ceRNAs identified during seed development

(A) Heatmap of ceRNAs. The color scale represents log_10_ (TPM+1) values. (B) The ceRNA number and total read counts of each stage. (B) The expression profiles of three ceRNAs and their parental genes.

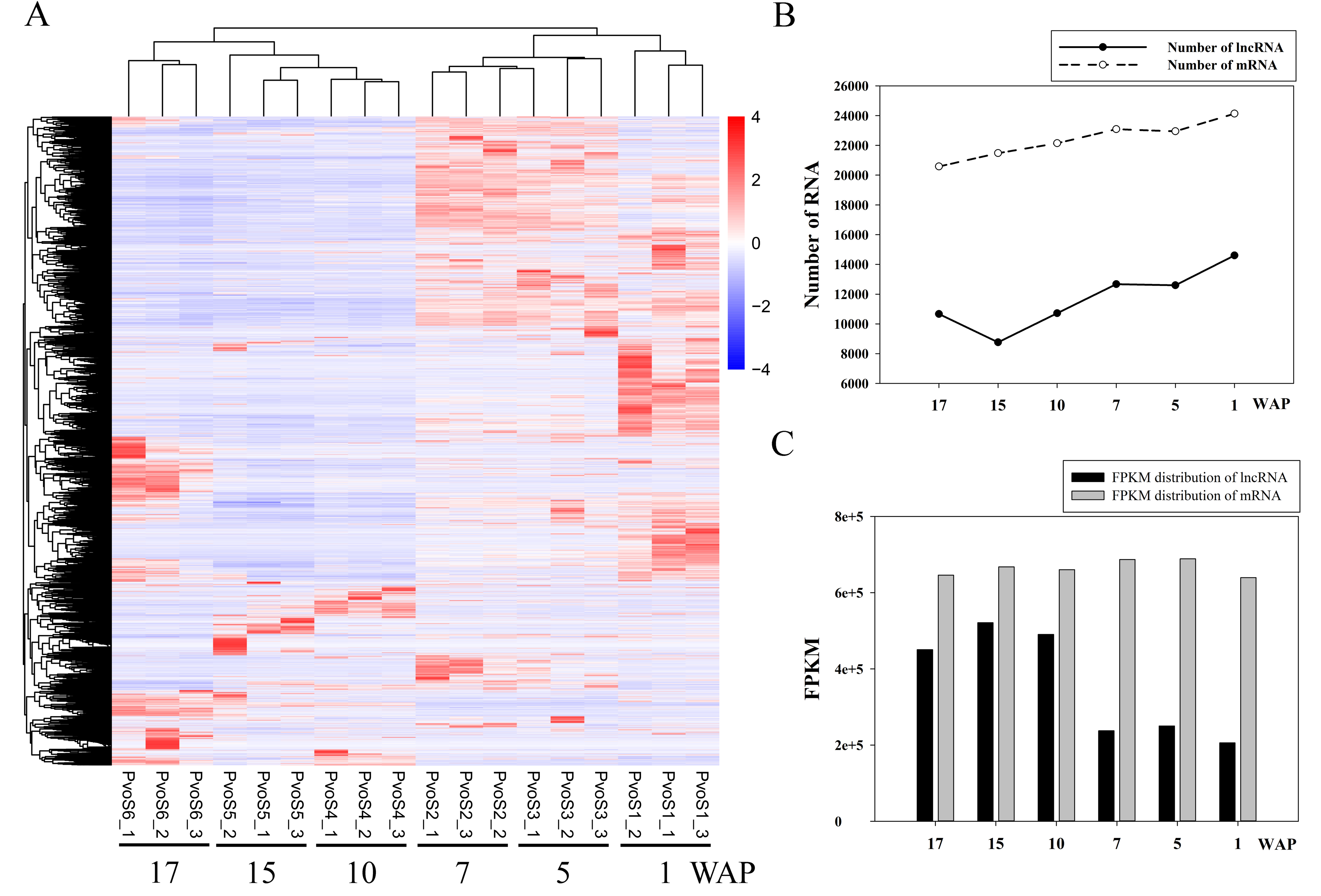

**Fig. S18** Summary of lncRNAs identified during seed development

(A) Heatmap of lncRNAs. The color scale represents log_10_ (RPKM+1) values. (B-C) The number (B) and FPKM distribution (C) of mRNAs and lncRNAs at each stage.
